# Integrated analysis of canine soft tissue sarcomas identifies recurrent mutations in *TP53, KMT* genes and *PDGFB* fusions

**DOI:** 10.1101/2023.01.06.522911

**Authors:** Sunetra Das, Rupa Idate, Susan E. Lana, Daniel P. Regan, Dawn L. Duval

## Abstract

Canine soft tissue sarcomas (STS) are a heterogenous group of malignant tumors arising from mesenchymal cells of soft tissues. This simplified collective of tumors most commonly arise from subcutaneous tissues, are treated similar clinically, and conventionally exclude other sarcomas with more definitive anatomical, histological, or biological features. Histologically, canine STS sub-types are difficult to discern at the light microscopic level due to their overlapping features. Thus, genomic, and transcriptomic profiling of canine STS may prove valuable in differentiating the diverse sub-types of mesenchymal neoplasms within this group. To this purpose we sought to characterize the transcript expression and genomic mutation profiles of canine STS. To delineate transcriptomic sub-types, hierarchical clustering was used to identify 4 groups with district expression profiles. Using the RNAseq data, we identified three samples carrying driver fusions of platelet derived growth factor B (*PDGFB*) and collagen genes. Sensitivity to imatinib was evaluated in a canine STS cell line also bearing a *PDGFB* fusion. Using whole exome sequencing, recurrent driver variants were identified in the cancer genes *KMT2D* (21% of the samples) and *TP53* (21%) along with copy number losses of RB1 and CDKN2A. Gene amplifications and resulting transcript increases were identified in genes on chromosomes 13, 14, and 36. A subset of STS was identified with high T-cell infiltration. This multi-omics approach has defined canine STS sub-types at a molecular level for comparison to their human counterparts, to improve diagnosis, and may provide additional targets for therapy.

## Introduction

Soft tissue sarcomas (STS) are a heterogenous group of mesenchymal neoplasms which represent approximately 8-12% of malignant tumors in dogs. Predominant within this category are hemangiosarcoma (HSA), fibrosarcoma (FSA), peripheral nerve sheath tumors (PNST), perivascular wall tumors (PWT) and histiocytic sarcomas, with myxosarcoma, liposarcoma, rhabdomyosarcoma, leiomyosarcoma, synovial cell sarcomas, and lymphangiosarcomas occurring less frequently. Approximately 6,000 to 25,000 cases are diagnosed in dogs yearly compared to approximately 12,500 STS cases diagnosed annually in adults and children in the United States (1). The subset of canine cutaneous and subcutaneous STS profiled here parallels current diagnostic and prognostic conventions in veterinary medicine (FSA, PNST, and PWT) and excludes those sarcomas with more definitive features (hemangiosarcoma, lymphangiosarcoma, histiocytic sarcoma, synovial sarcoma, leiomyosarcoma, and rhabdomyosarcoma). The FSA, PNST, and PWT tumors typically arise from subcutaneous tissues, are locally aggressive with low to moderate metastatic rates and are usually treated with surgical excision and/or radiation therapy (2). Previous studies have used immunohistochemistry, microarray, and RT-PCR techniques to identify markers to distinguish STS sub-types (3–5). Histologically, these STS sub-types can be difficult to discern due to overlapping morphology and their controversial cell of origin (6). Thus, genomic, and transcriptomic profiling of canine STS may prove valuable to differentiate these sub-types of mesenchymal neoplasms and understand their biological and clinical behavior.

To date a limited number of genome wide studies have been conducted in canine STS. Whole exome sequence analysis of tumor and matched normal samples from 10 dogs with STS (seven PNST, one FSA, one myxosarcoma, and one synovial cell sarcoma) identified single nucleotide substitutions in *NF1, MLL3,* and *PTCH1,* and amplification of *MDM4* in 4 separate tumors (7). The only recurrently mutated gene was *ATP7B* (n=2), while *AIG1* was amplified in 2 tumors. Another study identified recurrent changes in the TP53 pathway among 6 primary malignant PNST (8). In contrast, human PNST is strongly associated with neurofibromatosis type 1 with a 10-15% lifetime risk of PNST development (9). Recent next-generation sequencing of human malignant PNST identified recurrent mutations in NF1 (87.5%), TP53 (40%), and CDKN2A (75%) in addition to a dysregulated Ras pathway (10).

Canine FSA is frequently associated with rearrangements of chromosome 11 including loss of heterozygosity in the region containing CDKN2B-CDKN2A (11). In contrast, FSA in both children and adults are increasingly categorized based on frequently observed gene fusions. For example, congenital FSA often bears ETV6-NTRK3 gene fusions that activate the Ras-MAP kinase and PI3 kinase signaling cascades (12). The majority of adult human low-grade fibromyxoid sarcomas have FUS-CREB3L2 or FUS-CREB3L1 fusions, with the highly aggressive sclerosing epithelioid fibrosarcomas prevalently bearing EWSR1-CREB3L1 fusions (13). In addition, greater than 90% of Dermatofibrosarcoma Protuberans (DP) tumors carry COL1A1-PDGFB fusions leading to both improved diagnosis and neoadjuvant therapy with PDGFR inhibitors for patients with recurrent, nonresectable, or metastatic disease (14).

In this study, we have utilized a multi-omics strategy to delineate transcriptomic, genomic, and immunogenomic profiles of canine STS tumors. These 29 STS tumors identified histologically as FSA and PNST, were grouped by gene expression profiles into a single PNST cluster and 3 FSA clusters. The genomic profile identified *TP53* and *KMT2D* as the top two recurrently mutated genes with other identified driver mutations in cell cycle, DNA repair, and chromatin remodeling pathways. One of the major findings was identification of *PDGFB* fusions with collagen genes in 10% of the STS tumors. Both immunohistochemical and genomic immune profiling identified a set of tumors with high infiltration of immune cells. Taken together, exploration of these STS tumors reveals distinct groups of tumors with discernable genomic, transcriptomic, and immune-genomic profiles.

## Materials and Methods

### Canine tumor sample preparations

Soft tissue sarcoma (STS) tumors (n=29) were collected along with matched normal stroma (n=29) prior to treatment and stored at −80°C. Following the manufacturer’s protocol for TRIzol (Invitrogen, Catalogue # 15596026), genomic DNA was extracted from tumors and matched normals and RNA was extracted from tumor samples only. The DNA and RNA samples were purified using DNeasy or QiaAMP DNA Blood mini kits and RNeasy (Qiagen, Catalogue #69504, 51104, 74004), respectively, quantified on a NanoDrop Microvolume Spectrophotometer and quality was assessed by TapeStation or Bioanalyzer (Agilent). The total RNA was processed using Universal Plus™ mRNA-Seq library preparation kits with NuQuant® (product number: 0520-A01) and the resulting cDNA library was sequenced on Illumina NovaSeq6000 to generate 150 bp paired end reads. The SureSelectXT Target Enrichment System for Illumina Paired-End Multiplexed Sequencing Library kit was used to create genomic DNA libraries that were also sequenced on a NovaSeq6000 generating 150 bp paired-end reads.

### RNAseq pipeline

Low quality reads and adaptors were eliminated from Illumina RNAseq reads with Trimmomatic(15). The paired end reads (45.8 - 64.5 million) were mapped against CanFam3.1 with STAR (16), quantified using HTSeq-count (17), normalized with DESeq2 median of ratios method (18), and log-transformed for plotting and downstream analyses. STAR aligner mapped 94% of trimmed reads to CanFam3.1 genome. Unsupervised hierarchical clustering grouped samples into 4 optimal clusters which was based on the elbow method (19) using 1,926 genes with mean log2 expression >2 and mean variance >5. Spearman correlation and Ward’s minimum variance method was used to calculate distance and cluster STS samples, respectively. Gene fusions were identified via STAR-fusion by using trimmed fastq reads as input and curated to identify fusions with known cancer genes (20).

The differentially expressed genes (DEGs) between samples from each cluster and combined samples from the other clusters (e.g. H1 vs H2,H3,H4) were identified using DESeq2 (18) and functionally annotated with the g:Profiler R package (21). The pathways with FDR of <0.05 were considered enriched pathways for each cluster. For details see Supplementary Methods.

### Whole exome sequencing pipeline

The Illumina reads were processed to eliminate low quality reads and adapters using Trimmomatic and mapped against CanFam3.1 genome (https://may2021.archive.ensembl.org/Canis_lupus_familiaris/Info/Index) via BWA (22). Duplicate reads were marked with Picard and somatic variants were called using Mutect2 (23). Germline variants were eliminated using a panel of normals created from 118 normal samples following the GATK pipeline. Additionally, 90 million population variants, called from 722 dogs, were used as the germline resource option within Mutect2 when calling somatic variants (24). The remaining variants were processed using the filterMutectCalls GATK function and variants with a PASS notation in the FILTER column were characterized as somatic variants. The simple somatic (single nucleotide – SNV, and insertions and deletions – INDEL) variants were annotated using Ensembl Variant Effect Predictor (VEP, v99). The mutational signatures of each sample were deduced by using the MutationalPatterns R package (25).

The variants within cancer genes, as curated in COSMIC and OncoKB databases (26) were extracted from the list of somatic variants and putative drivers were identified as variants homologous to human cancer gene variants considered oncogenic/likely oncogenic. Pathway analysis of the mutated genes was conducted using g:profiler R package using hypergeometric tests to determine significant pathway/gene ontology terms.

The allele-specific copy number variant (CNV) genes were identified using Sequenza (27). Allele frequencies were used to calculate tumor normal depth ratio, allele-specific segmentation using the copynumber R package. The resulting segmentation file was annotated and genes with recurrent CNVs were identified with GISTIC2.0 (q-value ≤0.05). Genes with amplitude changes significantly correlated with transcript expression levels (Pearson correlation) were identified and cancer genes (OncoKB) were selected for plotting and cross-species comparison.

### Drug Sensitivity Assays

Canine cell lines (C2, STSA-1, and FACC-19CSTS36) were plated on 96-well plates at 2,000 cells/well in DMEM with 10% FBS. Following adherence for 24 hours, cells were treated with serial dilutions of imatinib (Gleevec, 100 μM to 1.7 nM, 3-5 wells/concentration) or DMSO vehicle. After 72 hours, cell confluence was monitored on the IncuCyte ZOOM Live-Cell Analysis System (Essen BioScience). Cell number was normalized to time zero readings and fit to 4 parameter non-linear curves of log dose versus percent of control and LD50 values were interpolated as the dose at which cell number was 50% of control.

### Immunogenomic profile

Scores for 22 of immune cell types were extracted and deconvoluted using CIBERSORT to cross reference against immunohistochemical staining of tumor-infiltrating CD3+ T-cells (28). Archived, formalin-fixed, paraffin embedded (FFPE) tumor samples were obtained from either the Colorado State University Flint Animal Cancer Center Tissue Archive or Veterinary Diagnostic Laboratory and were sectioned at five microns for H&E staining and immunohistochemical labeling. Immunolabeling was performed via routine, automated methods on the Leica Bond Max autostainer using a mouse monoclonal anti-human CD3 antibody (Leica, clone LN10). Whole slide images of IHC stained slides were digitally captured using an Olympus IX83 microscope at 10x magnification and fixed exposure times. Quantitative image analysis was performed using ImageJ software as previously described (29). The number of infiltrating immune cells was expressed as total positive cells per mm^2^ of tumor tissue area. See Supplementary Methods for details.

### Statistical analysis

All statistical analysis were done using R statistical tool. Significantly different groups of tumors with T-cell infiltration were analyzed using non-parametric Kruskal-Wallis test. For correlation analysis, Pearson correlation coefficient were calculated and FDRtool R package was used for correcting multiple testing of p-values.

### Data availability

The tumor and matched normal Illumina reads from WES were submitted to SRA database under Bioproject ID PRJNA882864. The RNAseq sequencing reads as well as raw count data for CanFam3.1 genes from 29 samples were submitted to GEO database (accession number: GSE214154). The list of commands, tools, R packages and their versions is available at https://github.com/sdas2019/Canine-soft-tissue-sarcoma-omics-pipeline.

## Results

### Canine patients and tumor overview

For this multi-omics study, STS tumors excised from 29 canine patients were reviewed by a veterinary pathologist and diagnosed into two major histologic sub-types of STS based on morphology alone: fibrosarcoma (FSA, n=15) and peripheral nerve sheath tumor\perivascular wall tumor (PNST n=14) (Table 1). While we recognize that the latter subtype represents a single morphological entity whose cellular origin remains controversial, investigating this distinction was not a primary goal of this study. Most dogs in this study were mixed breed (n=11) with Labrador retrievers (n=5) representing the single largest breed group and 41% of the 29 patients were female. Twenty-four patients had sub-cutaneous tumors of the extremities, with the remaining tumors in the oral cavity. Of the tumors selected for this study, (44%) were grade III, with increased potential for recurrence and metastasis (30). Histologically, FSAs were characterized by proliferations of spindle shaped cells arranged in streaming interwoven bundles or herringbone patterns in a background of dense collagenous stroma, and a lack of whorling. A subset of 7 FSA samples (T1124, T1238, T1292, T1407, T1490, T1834, and T416) were poorly differentiated\anaplastic, characterized by sheets of neoplastic cells with marked cellular and nuclear pleomorphism, polygonal to ovoid cellular morphology and containing large vesicular nuclei. In contrast, tumors diagnosed as PNST/PWT were characterized spindle shaped cells arranged in whorling, and storiform or palisading (Antoni A) growth patterns, or loosely arranged hypocellular regions within a myxoid matrix (Antoni B). No definitive histological features suggesting origination from vessels or nerves (besides whorling around central unidentifiable structures) was observed. However, to determine whether gene expression could further distinguish the tumors histologically identified as PNST/PWTs, the expression of pericyte markers was extracted (31). No significant up-regulation of these 17 genes were observed (Supplementary Fig. S1), while many samples demonstrated enrichment of genes and pathways consistent with peripheral nerve sheath origin, as detailed below.

**Table 1.** Patient and tumor characteristics.

| Sample ID | Breed | Sex | Age at diagnosis | Tumor sub-type | Tumor Location | Tumor Grade | Gene expression cluster |
| --- | --- | --- | --- | --- | --- | --- | --- |
| T1860 | Cock-a-poo | Male | 13.5 | PNST | Neck | II | H1 |
| T1921 | Golden Retriever | Female | 11.1 | PNST | Flank | I | H1 |
| T2063 | Mix | Female | 10.2 | PNST | Thigh | II | H1 |
| T148 | Mix | Male | 14.8 | PNST | Hock | I | H1 |
| T218 | Mix | Female | 11.5 | PNST | Hip | II | H1 |
| T230 | Mix | Female | 13.3 | PNST | Knee region | I | H1 |
| T311 | Mix | Male | 7.5 | PNST | Inguinal region | I | H1 |
| T320 | Mix | Male | 15.5 | PNST | Flank | II | H1 |
| T336 | Mix | Female | 11.3 | PNST | Rib region | II | H1 |
| T344 | Basset Hound | Female | 10.8 | PNST | Humerus | I | H1 |
| T697 | Labrador Retriever | Female | 8.9 | PNST | Thorax | III | H1 |
| T1238 | Labrador Retriever | Male | 9.1 | FSA | Hock | III | H2 |
| T167 | Mix | Male | 8.6 | FSA | Cranial sternum | III | H2 |
| T1891 | Labrador Retriever | Male | 9.7 | FSA | Femur | II | H2 |
| T1989 | Labrador Retriever | Female | 11.1 | FSA | Neck | II | H2 |
| T416 | Alaskan Malamute | Male | 10.2 | FSA | Elbow joint | III | H2 |
| T479 | German Shepherd | Female | 9.3 | PNST | Hip | II | H2 |
| T738 | Pointer | Male | 5.8 | PNST | Tarsus | III | H2 |
| T1124 | Greyhound | Male | 8.3 | FSA | Side, lateral | III | H3 |
| T1407 | Mix | Male | 3.9 | FSA | Maxilla | III | H3 |
| T1490 | Golden Retriever | Male | 5.4 | FSA | Hip | III | H3 |
| T1834 | Mix | Male | 12.6 | FSA | Mandible | III | H3 |
| T232 | Poodle | Male | 6.9 | PNST | Femur | III | H3 |
| T1292 | Beagle | Male | 12.4 | FSA | Maxilla | III | H4 |
| T1462 | Labrador Retriever | Male | 11.9 | FSA | Thigh | III | H4 |
| T1585 | Beagle | Female | 10.0 | FSA | Abdomen | III | H4 |
| T1923 | Siberian Husky | Male | 10.5 | FSA | Rib region | II | H4 |
| T282 | Border Collie | Female | 12.5 | FSA | Lip | II | H4 |
| T485 | Mix | Female | 6.3 | FSA | Maxilla | I | H4 |
| PNST: Peripheral nerve sheath tumor; FSA: Fibrosarcoma |  |  |  |  |  |  |  |

### Transcriptomic profiling of STS

#### Gene expression defined tumor clusters

The RNAseq reads ranging from 45.8 - 64.5 million were trimmed and 94% of these reads were mapped against CanFam3.1 genome to obtained gene expression data (Supplementary Table S1). Unsupervised hierarchical clustering of 1,926 gene grouped 15 FSA and 14 PNST samples into 4 optimal hierarchical (H) clusters with 11, 7, 5, and 6 samples in H1, H2, H3, and H4 clusters, respectively (**Fig. 1,** Supplementary Figure S2) (19). Comparison of histological categorization and tumor clustering revealed that all tumors in cluster H1 were PNSTs. However, three samples histologically identified as PNSTs were placed in cluster H2 (T738, T479) or H3 (T232) (Fig. 1A). The multi-dimensional scaling plot shows that clusters H1 and H2 are separated while H3 and H4 display some overlap (Fig. 1B).

**Fig. 1.**
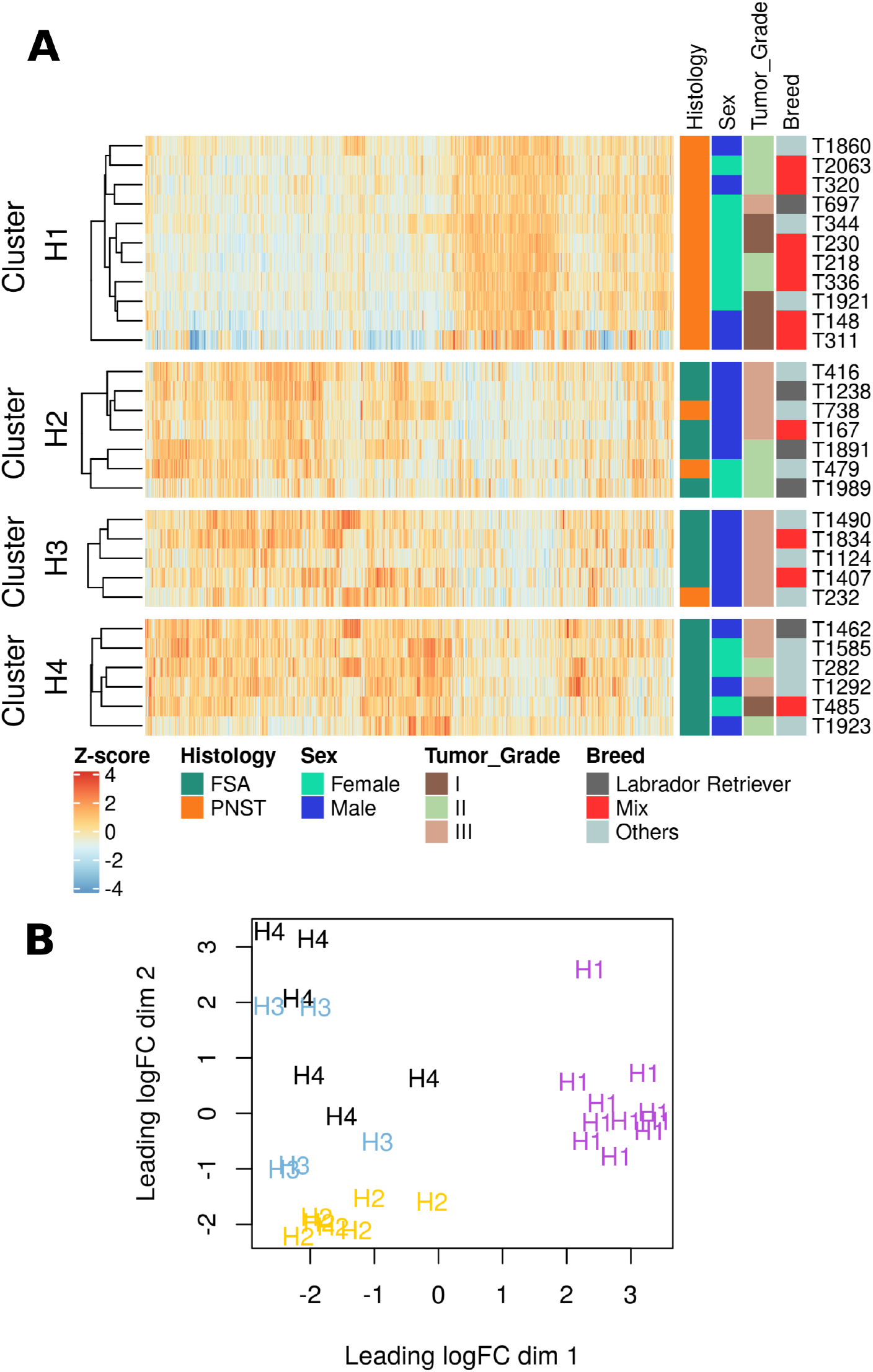
Clustering of canine soft tissue sarcoma samples. A. Unsupervised hierarchical clustering of 29 STS tumor samples using genes with variance >log2 5 and mean >log2 2. The samples were clustered using the spearman correlation distance method and grouped into 4 clusters. The clinical features associated with samples were plotted on the right side of the heatmap. B. Multidimensional scaling (MDS) plot visualizing principal component analysis in two dimensions. Tumor samples with similar gene expression profiles were clustered together.

Comparisons were made between samples from one cluster and combined samples from the remaining clusters to identify the genes and pathways distinguishing these 4 clusters. The significantly up-regulated genes (H1=575, H2=94, H3=119, H4=430, Supplementary Table S2) in each cluster were analyzed to identify pathways and gene ontologies driving the four clusters (Supplementary Table S3). The top 10 upregulated genes in each of the 4 clusters are plotted in Fig. 2A. The biological themes identified through pathway analysis for H1, H2, H3, and H4 clusters were associated with nervous system-, extracellular matrix-, development-, and immune-related genes, respectively (Fig. 2B). The heatmap of selected genes significantly upregulated in H1 samples involved in neuronal cell body (GO:0043025), neurogenesis (GO:0022008) and neuron differentiation (GO:0030182), includes *GAL, CHRNA4, NRXN1, MAST1, ROBO1, TNR, CBLN1, ACTL6B, CRMP, NEFH,* and *NYAP.* Upregulated extracellular matrix (GO:0031012) genes in H2 samples include: *COL11A1, COL24A1, COL2A1, EMILIN3, SFRP1,* and *VIT.* In H3, the upregulated genes *BMPR1B*, *DNAH*, *HOXD11*, *SIX1*, *SIX2*, NPNT, WNT7A, were associated with pattern specification (GO:0007389) and tissue morphogenesis (GO:0048729). In cluster H4, a significant proportion of enriched pathways were immune system related and included genes like *CCL3, CD27, CD79B, CD8A, CTLA4, IL21R, IL23R, and IL7R,* among others (Supplementary Table S3 and Supplementary Figure S3).

**Fig. 2.**
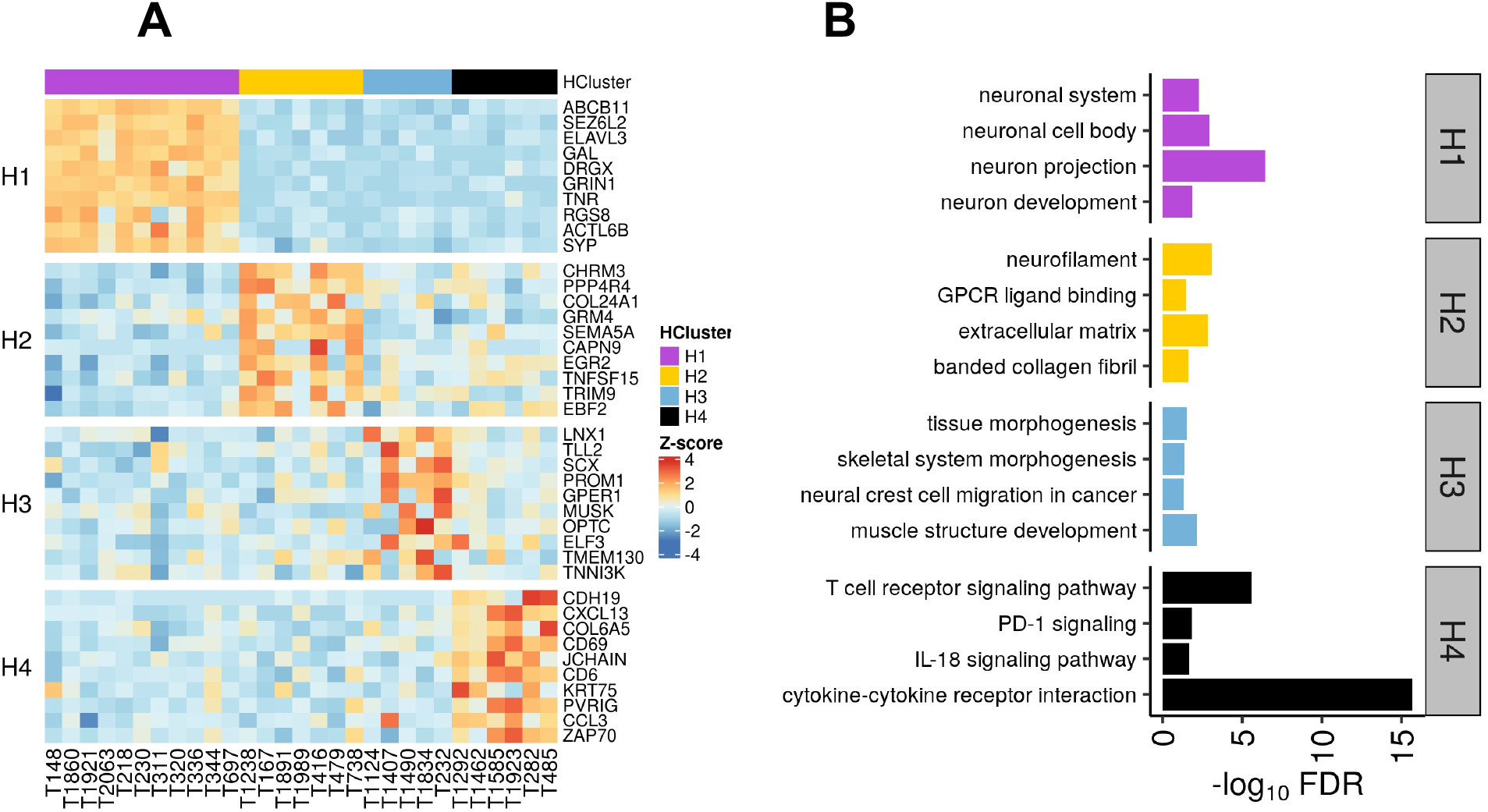
Enriched genes and pathways in 4 clusters A. Heatmap of top 10 significant differentially expressed genes (positive log2 fold change >2) in each cluster. B. Graphical representation of false discovery rate of representative pathways and gene ontologies identified by functional annotation of DEGs in each cluster. The main biological theme in cluster H1, H2, H3, and H4 were nervous system, extracellular matrix, development, and immune system-related, respectively.

### Genomic profile of STS

#### Mapping and variant calling statistics

Whole exome sequencing of 29 primary STS tumors and corresponding matched normals was mapped against the CanFam3.1 genome and somatic short variants and copy number variants (CNVs) were identified. Following trimming, 90.7% (±0.44%) reads were retained and 99.9% (±0.0003%) of those reads were mapped to the CanFam3.1 genome (Supplementary Table S4). The average coverage of mapping was 95.9X (±28.3X) and ranged from 22.5X (T282) to 132.7X (T1238) (Supplementary Fig. S4).

#### Putative mutational signatures of canine STS

The distribution of 6 types of single nucleotide changes shows that C < T was most prevalent, followed by C<A (Supplementary Fig. S5A). Non-negative factorization (NMF) generated three *de novo* signatures that were assigned to the tumors and named based on their similarity to human COSMIC signatures. The SBS5-like signature, (etiology: correlated with age of patients) was assigned to 52% of the tumors, 38% of the tumors were SBS6-like (etiology: defective DNA mismatch repair), and 10 % were considered SBS29-like (etiology: tobacco chewing habit; Fig. 3, Supplementary Fig. 5B) (32). There was no difference in signature distribution across the 4 clusters of tumors, however, the SBS29-like signature was restricted to clusters H3 and H4. The estimation of cosine similarities between individual tumor mutational matrices and 60 COSMIC signatures showed that in addition to SBS5 (n=14), SBS6 (n=3) and SBS29 (n=1), some samples had greater similarity to SBS3 (defective homologous recombination-based DNA damage repair, n=3), SBS30 (deficiency in base excision repair due to inactivating mutations in NTHL1, n=2), SBS40 (etiology: unknown, n=5) and SBS94 (etiology: unknown, n=1) (Fig. 3C).

**Fig. 3.**
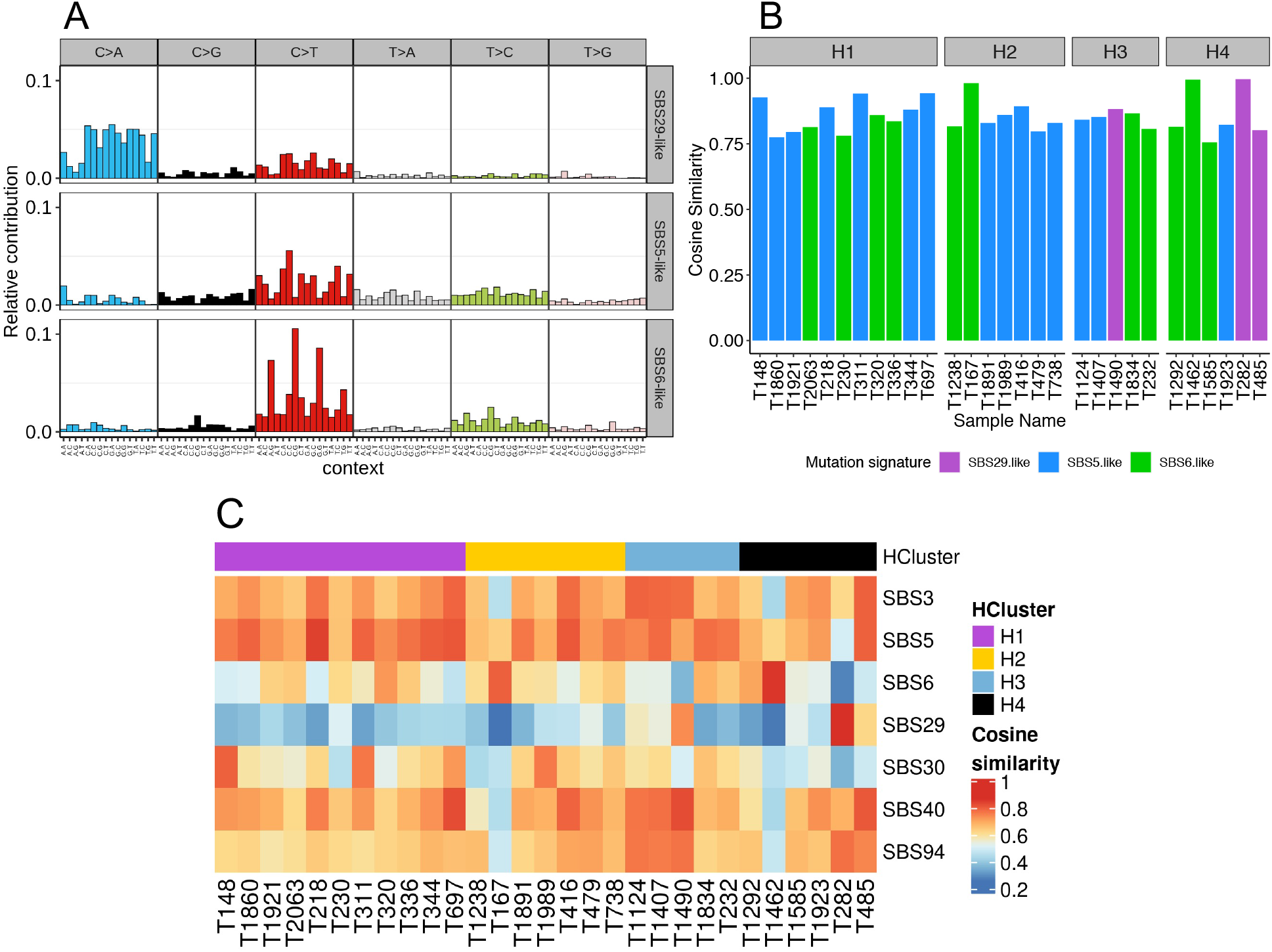
COSMIC mutational signature analysis in canine STS tumors. A. Relative contribution of 96 trinucleotide contexts across three predominant *de novo* signatures as identified by non-negative matrix factorization method of SNV frequencies from samples. The three *de novo* signatures were similar to COSMIC signatures SBS5, SBS6, and SBS29. B. Distribution of *de novo* signatures across 29 STS samples. C. Heatmap of cosine similarities between COSMIC mutational signature profiles and the single nucleotide mutation frequency profiles of 29 STS tumors. For each sample, the signature with the highest cosine similarity was chosen to plot. This resulted in the selection of 7 COSMIC mutational signatures.

### Mutations in STS tumors

#### SNVs and INDELs

The number of SNVs ranged from 193 (T2063) to 3,744 (T282) and INDELs range was 43 (T1238) to 1,281 (T311) (Supplementary Fig. S6). A total of 6,439 non-synonymous coding sequence (CDS) variants in 4,254 genes were extracted from 29 samples ranging from 13 (T1238) to 1,329 (T282) (Supplementary Table S5). The top recurrently mutated genes, present in 21% of samples, included *KMT2D, TP53, SAFB, ZFN398, TTN,* OR13L2, and COR52A16 (Supplementary Figure S7). The median purity of tumors was estimated to be 0.54 (range: 0.31 – 1), although, there was no correlation between purity and number of variants identified (Pearson’s correlation coefficient: 0.16, p-val=0.39).

The median protein coding mutations per callable megabase (coverage >10X) was 2.7 and ranged from 0.03 (T1238) to 37.3 (T282) (Fig. 3). Missense mutations were most prevalent (median 62% of all CDS mutations), followed by frameshift and UTR regions variants (13% each) (Supplementary Figure S8). Using the SIFT scoring method, the identified deleterious missense mutations ranged from 2 in T1238 to 553 in T282 with a median of 29 across all tumor samples. Functional annotation of CDS variant genes identified 116 pathways (FDR <0.05, Supplementary Table S6). Relevant pathways included: PI3K-Akt signaling (72 % tumors), focal adhesion, RHO GTPase cycle (58 % tumors), signaling by PDGF (52% tumors) and chromatin binding (83% tumors). Since most somatic variants likely represent passenger mutations, we identified putative driver genes by cross-referencing canine somatic variants to likely human oncogenic variants. About 6.6% (n=427) of CDS variants were within OnkoKB designated cancer genes with 55 variants in 37 genes identified as putative drivers (26). The median number of cancer genes in each tumor was 9 (range 0 – 75). These genes were grouped into 7 biological categories: cell cycle and DNA repair, chromatin organization/binding, PI3K-Akt-mTOR-signaling pathway, Wnt signaling pathway, transcription regulator activity, nervous system development, and kinase binding (Fig. 4). The bulk of the putative driver variants (37%) were in chromatin organization/binding genes, followed by 18% categorized in the cell cycle and DNA repair and nervous system development gene groups. About 28% of samples carried mutations in *KMT* genes, most prevalent was *KMT2D* mutated in 6 samples: T311, T167, T416, T282, T1462, T1891 (Supplementary Figure S9). There were 10 STS tumors with alterations in cell cycle and DNA repair genes that included recurrently mutated *TP53* (5 samples: T1124, T1490, T336, T416, T1834, T1462). Among the 8 putative driver genes grouped in nervous system development, five genes were mutated in tumors that were histologically identified as PNSTs.

**Fig. 4.**
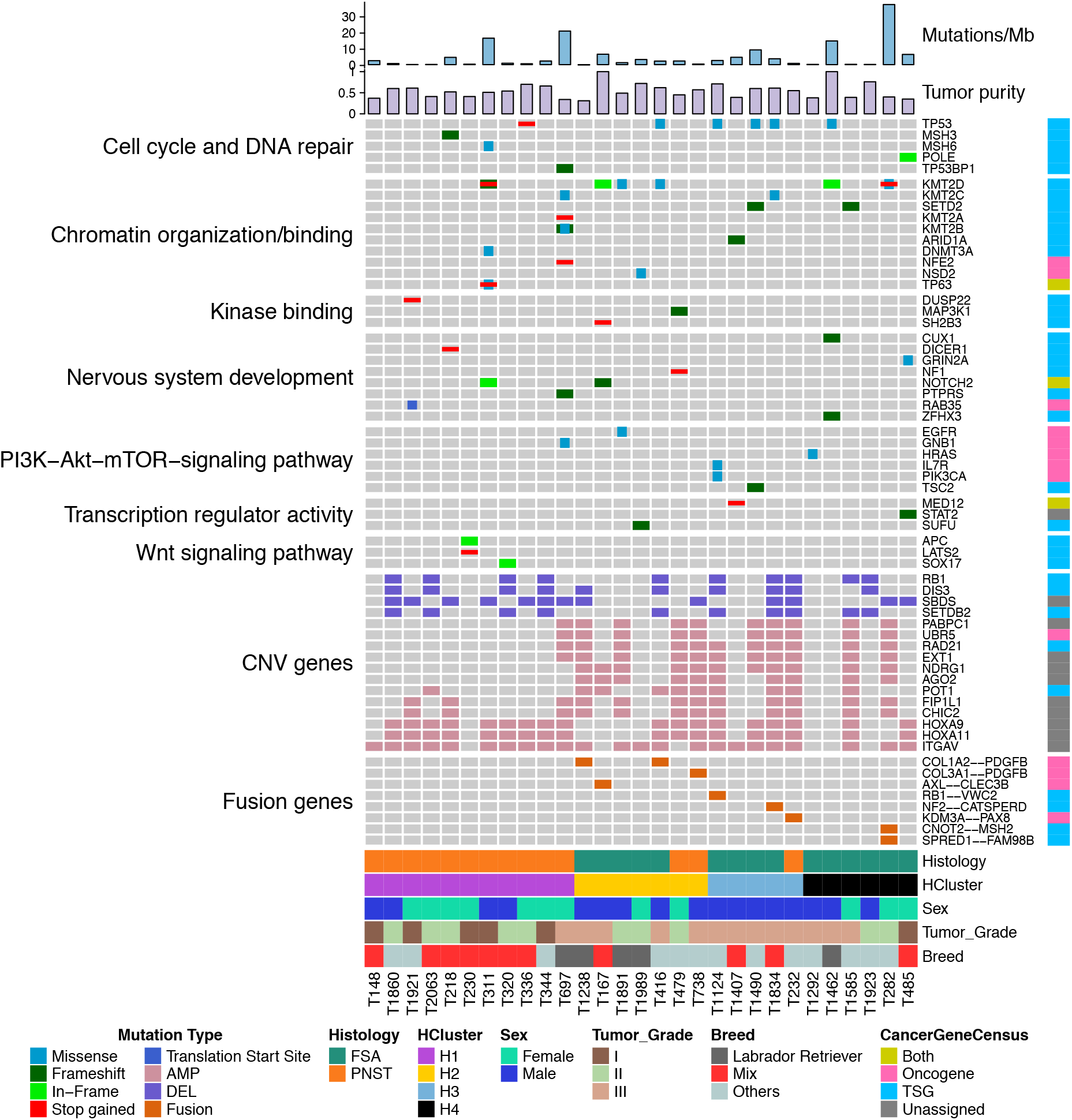
Mutational landscape of canine STS. Oncoplot of selected putative cancer driving variants. INDELS or SNVs are arranged in biological pathways, cancer genes with copy number variations significantly correlated to gene expression (CNV; AMP: amplifications, DEL: deletions) and fusion genes are illustrated. The clinical parameters for each of the 29 samples were represented in the bottom of the Oncoplot, along *in silico* defined cluster ID. The cancer genes were categorized into oncogene, tumor suppressor gene (TSG), both or unassigned as defined by OncoKB database. Additionally, the top two bar plots included mutational burden scores and tumor purity values per sample.

#### Copy number variants

We identified 1,203 genes that were located within significantly amplified or deleted regions of canine genome (Supplementary Table S7). The median number of altered copy number variation (CNV) genes across 29 samples was 576 (range 8 (T230) to 1093 (T1834) Supplementary Fig. S10). There were significantly more amplified genes (77.6%) (Student T-test, p-value 1.3e-07). The only significantly enriched functional pathway was the Hippo signaling pathway, and enriched gene ontologies included blood vessel morphogenesis possibly associated with angiogenesis (Supplementary Table 8). Cancer genes, as designated in the OncoKB database, were 5% of all CNV genes with 51 and 13 significantly amplified and deleted genes, respectively. To evaluate the functional effect of these CNVs, we used Pearson correlations between CNV gene amplitude values and Z-scored gene expression values. This identified 242 amplified and 68 deleted genes positively correlated with corresponding expression levels (Supplementary Table 9). Of these, 16 were known cancer genes restricted to chromosomes 6, 22, 13, 14 and 36. This included 8 amplified genes from CFA13: *AGO2, PABPC1, UBR5* (each altered in 35% of tumors), *RAD21, NDRG1, EXT1* (38%), *FIP1L1, CHIC2* (41%); 3 CFA14 genes: POT1 (35%), *HOXA9, HOXA11* (62%), and *ITGAV*(CFA36, 79%). Of the 4 deleted genes: *DIS3, RB1, SETDB2* (35%), were on CFA22, while *SBDS* (45%) was on CFA6 (Fig.4). The CFA13 amplified genes were largely restricted to clusters H2-H4 except for T697.

#### Fusion variants

Using the RNAseq data, 234 unique fusion genes were identified, where 121 were interchromosomal and 113 were intrachromosomal fusions. To identify putative drivers, fusions with gene partners from the same gene family and genes without annotations were eliminated and the remaining fusions were cross referenced against the Cbioportal database to identify likely oncogenic changes due to loss of TSG or activation of oncogenes. This identified 15 putative oncogenic fusions within 9 samples (Supplementary Table S10) and a subset of these are depicted in Fig. 4. Most of these putative driver fusions (n=12) were identified in clusters H2 and H3, with a single fusion (CAND2--PPARG) in H1 and two (CNOT2--MSH2, SPRED1--FAM98B) in the H4 clusters.

Three tumors had in-frame fusions of collagen genes (exon 6 *COL1A2,* exon 20 *COL1A2* and exon 35 *COL3A1*) with exon 2 of platelet derived growth factor subunit B (*PDGFB*) which were confirmed by Sanger sequencing of amplified cDNA (T1238/T416 and T738, respectively, Supplementary Figure S11). These fusions were associated with significantly increased expression of *PDGFB* (Z-score: 4.22 in T416, 2.16 in T738, 1.11 in T1238) in these tumors compared to the cohort (Supplementary Figure S12). Of these 3 tumors, T1238 and T738 also have reduced *CDKN2A* expression (Z-score: −0.69, −0.65), while tumor T416 bears a TP53 missense mutation and *RB1* CNV loss (Supplementary Figure S13). These 3 tumors were in the H2 FSA cluster associated with extracellular matrix and G-protein coupled receptor signaling pathways.

Similarly, increased *EGR2* expression in T1238, T167, and TT416 was associated with EGR2--ATXN7 fusions. Perhaps more relevantly, we also observed loss of TSG transcripts associated with NF2 fusions in T1834, a CD81--MEN1 fusion in T1238, and reduced RB1 from the RB1--VWC2 fusion in T1124 (Fig. 4). While the CNOT--MSH2 fusion in T282 was associated with elevated *MSH2* transcripts, the resulting frameshift in the MSH2 DNA mismatch repair gene may produce nonfunctional protein. In fact, T282 carried the highest mutational burden of any tumor in this study with 37.63 mutations/Mb.

A newly developed canine soft tissue sarcoma cell line, FACC-19CSTS36, with a Sanger sequence confirmed *COL3A1-PDGFB* fusion was tested for sensitivity to imatinib, a tyrosine kinase inhibitor capable of inhibiting v-Abl, c-Kit, and PDGFR. Also included in this analysis as positive and negative controls were the canine C2 mast cell line, with an internal tandem duplication that activates c-Kit, and the canine STSA-1 soft tissue sarcoma cell line with an inactivating frameshift mutation in NF-1 (33). The C2 cell line, as expected, was sensitive with an interpolated LD50 of 24 nM, while LD50 values for the STSA-1 and FACC-19CSTS36 cells were approximately 1000-fold higher at 17 and 45 μM, respectively (Supplementary Figure S14).

#### Markers for FSA and PNST

Previous studies have identified 15 and 7 marker genes for PNST and FSA. respectively (4,5). We confirmed significantly elevated expression of 7 of the 15 genes in PNST samples (n=14): *CLEC3B, GLI1, FMN2, NMUR2, DOK4, HMG20B* and *GAP43* (Fig. 5A). The remainder of the marker genes were either significantly down-regulated in PNST samples (*NES, ROBO1, NGFR*) or unchanged (*EGR2, MBP, MPZ, KIF1B, SOX10,* Supplementary Figure S15).

**Fig. 5.**
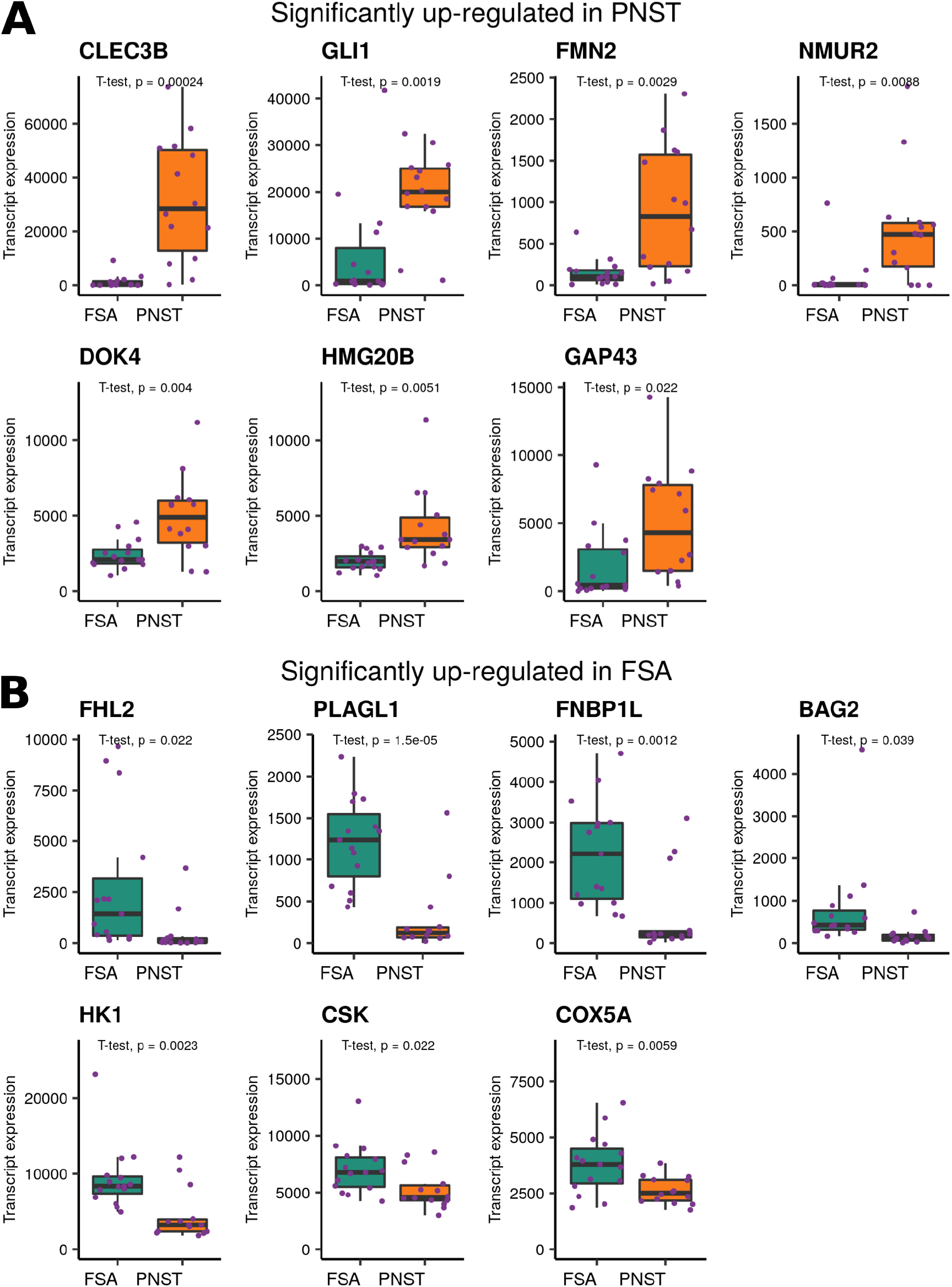
Expression levels of previously identified PNST and FSA marker genes. A. Transcript expression boxplots of seven previously identified PNST marker genes. These genes had significantly higher transcript expression in 14 canine PNST samples when compared to 15 FSA samples. B. Transcript expression boxplots of previously identified FSA marker genes with significantly higher expression in FSA samples in comparison to PNST samples. The bar in boxplots represents median transcript expression. Expression levels plotted here are normalized count data.

The seven genes reported as markers from FSA samples were *FHL2*, *PLAGLI*, *FNBP1L*, *BAG2*, *HK1, CSK* and *COX5A* (4). Parallel to the previous publication, mean expression of these genes was significantly higher in FSA samples (n=15) (Fig. 5B).

#### Immunogenomic profiling

The RNAseq transcriptome data was used to generate an immunogenomic profile of the STS tumors and compared to immunohistochemical staining in STS tumors (n=27) with the pan-T cell marker CD3. The median percent tumor area staining positive for CD3+ T cells was 0.135%, ranging from 0.006 in T479 (H2 cluster) to 7.557% in T1923 (H4 cluster). There was a significant difference in CD3+ T cell infiltrates across the 4 clusters of tumors (Kruskal-Wallis test, p-value 0.024) with elevated staining in cluster H4 samples compared to H1 cluster samples (Benjamini-Hochberg adjusted p-value 0.022), but not the H2 and H3 clusters (Fig. 6A, Supplementary Figure S16). Using Pearson correlations, we identified 319 and 8 genes positively and negatively correlated with the CD3+ immune score, respectively (Supplementary Table S11). The eight genes negatively correlated to CD3 staining were: *LMNA, RGS12, KIF7, LPAR2, LRRC4B, OBSL1, NAT14,* and *MAMDC4.* Of the 319 genes positively correlated to the IHC immune score, *IDO1, TNFRSF14, BTLA, TIGIT, PDCD1LG2, TNFRSF4, PDCD1, ICOS, CTLA4* and *CD274* are actionable immune checkpoint molecule targets approved or under clinical evaluation in human immuno-oncology studies. Enriched KEGG pathways linked with these genes included Cytokine-cytokine receptor interaction, T cell receptor signaling, PD-L1 expression and PD-1 checkpoint pathway in cancer, and B cell receptor signaling (Supplementary Fig. S17).

**Fig.6.**
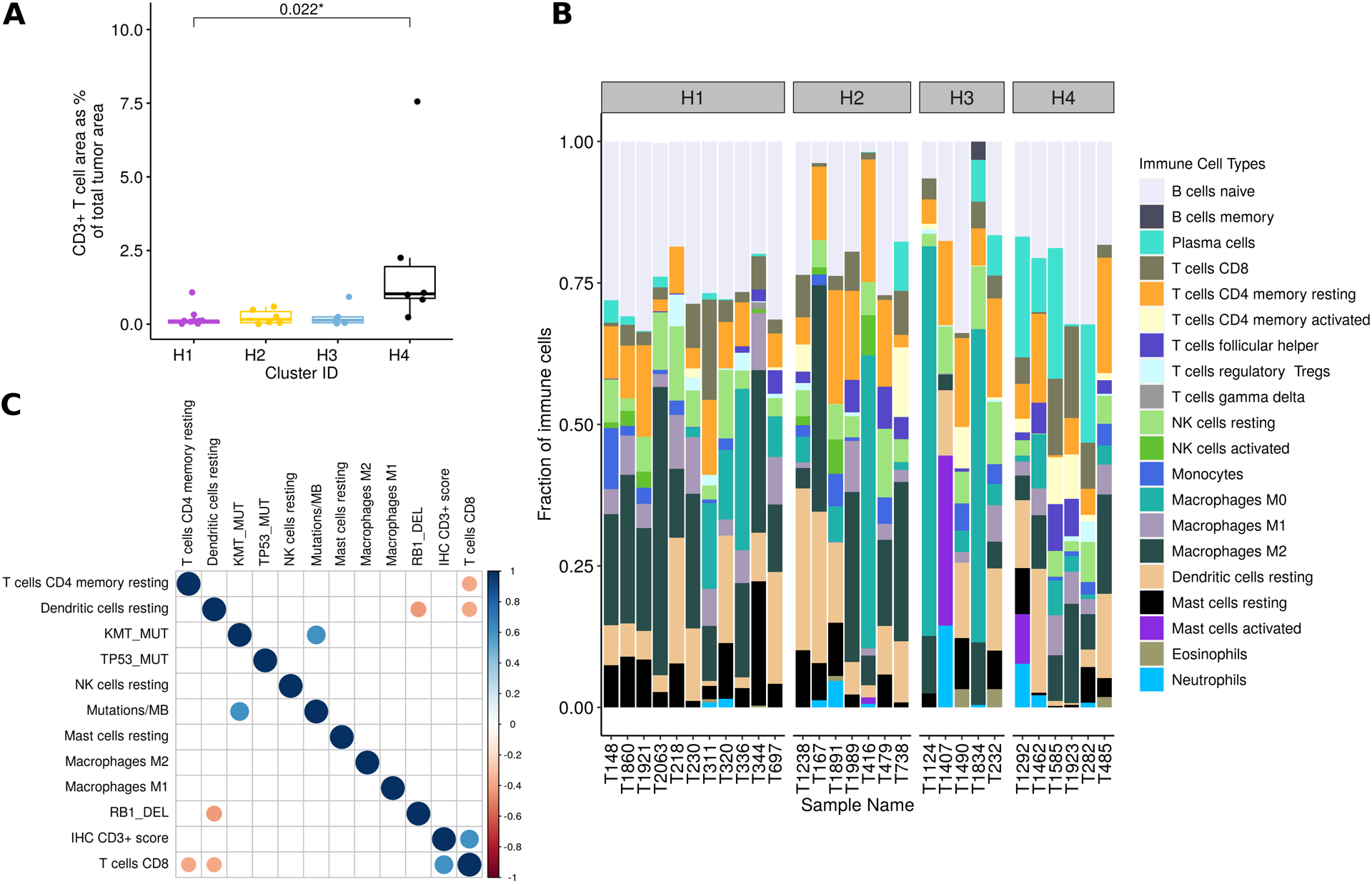
Immunogenomic profile of STS tumors. A. Barplot showing distribution of CD3+ T-cell density in each cluster, as determined by immunohistochemistry. The samples in the H4 cluster had a significantly higher density of intra-tumoral CD3+ T cells. B. Deconvolution of tumor microenvironment using CIBERSORT. The relative abundance of 20 cell types in each sample were plotted based on gene expression profiles. C. Correlation matrix of immune cell type scores generated using CIBERSORT, genomic variants, and immunohistochemical derived CD3+ T-cell density (IHC CD3+ score). Only the significant Pearson correlation coefficients were plotted. The CD3+ T cell IHC scores of 29 STS samples positively correlated with CIBERSORT relative frequencies of CD8 T-cells.

Additionally, CibersortX gene expression deconvolution method was used to characterize tumor infiltrating immune cells (28). The most abundant cell types infiltrating STS tumors were naïve B cells (n=12), macrophages M2 (n=5), macrophages M0 (n=4), dendritic cells resting (n=3), Plasma cells (n=2, T cells CD4 memory resting (n=2), and mast cells activated (n=1) (Fig. 6B). Additionally, consistent with CD3+ T cell immunostaining results, samples from cluster H4 had a higher fraction of CD8 T cells compared to samples in the remaining clusters.

To identify significant monotonic associations between the immune cell score, IHC score, mutations/MB, and genomic variables, Pearson rank correlation coefficients were computed (Fig. 6C). The genomic variables included samples with and without KMT gene mutations (KM_MUT), TP53 mutations (TP53_MUT, RB1 deletions (RB1_DEL). The CD3+ immune score positively correlated with gene expression scores for T cell CD8. There was an inverse correlation between the gene expression score for dendritic cell resting and scores for T cells CD8 and RB1_DEL. Increased mutational burden (mutations/MB) significantly correlated with *KMT_MUT* but not with any of the immune scores.

## Discussion

This study represents the first comprehensive genomic analysis of a cohort of 29 canine STS reporting on the whole exome and RNA-sequencing profiles of fibrosarcoma and peripheral nerve sheath tumor (PNST) subtypes. From the transcriptome analysis, we identified 4 clusters (H1 - H4) with distinct enriched biological features. We maintained this 4-cluster framework for downstream comparisons. Using WES and RNAseq gene fusion analysis we identified putative driver mutations associated with each of these STS subtypes, evaluated CNVs associated with transcript changes, and performed immune profiling of these clusters. Overall, this analysis begins to identify critical similarities and differences in the potential genetic drivers between these canine soft tissue sarcoma subtypes and their human counterparts.

Mutational signatures using relative frequencies of single nucleotide substitutions have been used to elucidate the underlying cancer etiology (32). Although mathematical association of mutational spectrum rates to various signatures is possible using WES data, mechanistic understanding of cancer causation needs experimental validation (34). In this study, a majority (52%) of canine STS exhibited the SBS5-like signature which is an age-related or clock-like human signature. In comparison, approximately 53% of human STS had signatures attributed to SBS5, while an additional 37% were attributed to another clock-like signature, SBS1 (35). In this human data set, 2 tumors with the highest mutational burden were associated with the SBS6 defective mismatch repair signature. Eleven of the 29 canine STS (38%) were designated with the SBS6-like signature. There was no difference in the age at diagnosis or TMB between the different signatures, nor was there a correlation between age at diagnosis and mutations/MB for the age-related signature SBS5. Of the 3 tumors bearing the tobacco chewing signature, SBS29 (high C>A substitutions), 2 were localized to the mouth, suggesting that a similar environmental exposure could be involved. Interestingly, both tumors were in the H4 cluster with elevated immune system-related genes which might link this signature to an inflammatory response.

We identified recurrent somatic mutations in *TP53* and *KMT2D* in 21% of the tumors. There was a positive correlation between mutations in KMT family genes and mutational burden, where *KMT2D* mutated tumors in the H1 cluster had two of the highest mutational rates. Loss of KMT2D function in human tumors has been associated with high mutational burden and immune checkpoint blockade response (36). Previously, Sanger sequencing of amplified exons 5-8 of TP53 revealed missense mutations in 20% of canine STS samples, with 3 of 7 variants identified in mPNST samples and no variants were identified among 3 fibrosarcomas tested (8). Southern blot analysis identified *MDM2* amplification in 3 of 7 PNST. The previous WES study of 10 STS tumors (7), including 7 PNSTs, did not identify recurrent cancer gene variants with MDM4 amplification in one tumor as the only TP53 pathway variant. Similarly, among the 14 histologically identified PNST tumors, we did not identify any recurrent mutations. An overall look at the P53/cell cycle pathway revealed that 66% of the STS tumors had dysregulation of *TP53, MDM2,* CDKN2A or *RB1* (Supplementary Fig. S12). Six tumors had variants in *TP53* with 5 missense mutations in FSAs, and a single PNST with a stop-gained variant. CNV evaluation identified *RB1* loss in 10 of the 29 tumors, but only modest reductions in transcript expression. Eight tumors had < 10% of average *CDKN2A* transcript levels; seven tumors were FSAs with 5 grouped in H2 cluster and significant *CDKN2A* homodeletion was identified in 4 FSAs. As seen in most human cancers, loss of *CDKN2A* transcript expression was generally exclusive of RB1 loss, while three tumors had both *TP53* variants and *RB1* loss and one tumor had *MDM2* overexpression with *RB1* loss (37). Finally, SBDS ribosome maturation factor (*SBDS*) CNV loss was identified in 45% of the tumors. Germline mutations in this gene lead to the Shwachman-Diamond syndrome and increased cancer incidence possibly due to SBDS regulation of TP53 (38). Of the 10 tumors without pathway modifications, 6 were in the PNST H1 cluster (54%), indicating that for this cluster other pathways are involved in pathogenesis.

Given the prevalence of gene fusions in subsets of human soft tissue sarcomas, we analyzed the RNAseq data to identify fusions that may play a role in tumor progression. Notable, among these fusions are in-frame fusions of the collagen genes (exon 6 *COL1A2,* exon 20 *COL1A2* and exon 35 *COL3A1*) with exon 2 of *PDGFB* in tumors of the H2 fibrosarcoma cluster which was associated with extracellular matrix and G-protein coupled receptor signaling pathways. Similar fusions of COL1A1-PDGFB have been identified in two human STS subtypes: dermatofibrosarcoma protuberans (DFSP) and giant cell fibroblastoma; and increased expression of *PDGFB* is considered predictive of therapeutic response to PDFGR inhibitors (14), although resistance subsequently develops (39,40). However, we found that cultured STS cells bearing a *COL3A1-PDGFB* fusion were only responsive to the PDGFR inhibitor, imatinib, at concentrations in the μmolar range, similar to a cell line without the *PDGFB* fusion or other targets for this multi-target tyrosine kinase. Since canine PDGFRA/B are 96% and 89% identical to the human proteins, lack of response does not seem to be due to species specificity. In fact, similar results have been observed in cultures of human DFSP cells (41), suggesting that therapeutic responses in these tumors may rely on inhibition of tumor-stroma interactions (42). Finally, losses of *CDKN2A* in these cell lines or *CDKN2A*/*RB1* in these tumors in the H2 group may also contribute to tumor development and modulate drug responses.

Malignant PNST in humans often develops subsequent to neurofibromatosis type I in which inactivation of the NF1 tumor suppressor gene (TSG) results in activation of the MAP kinase signaling pathway (43). Consequently, MEK inhibitors have been approved for treatment of manifestations of this syndrome. In addition, a variety of other sporadic human tumors are also associated with inactivating mutations in NF1 including glioblastoma, lung adenocarcinoma, acute myeloid leukemia, and ovarian and breast cancers (44). However, only one *NF1* variant was identified in an H2 cluster sample histologically identified as a PNST. Similarly, the human Legius syndrome, caused by germline mutations in *SPRED1,* which can limit binding to *NF1,* has similar but milder neurofibromatosis type I symptoms (45) and an increased incidence in pediatric leukemias with elevated ERK signaling. Here we provide the first report of a *SPRED1* fusion gene in a canine FSA sample. *SPRED1* mutations are observed in about 2% of human cancers with *SPRED1* loss attributed as a driver in 37% of mucosal melanomas (46).

In a previous study, we utilized a MAP Kinase pathway activity (MPAS) score developed to predict MEK1/2 inhibitor sensitivity in human cancers across a panel of canine cancer cell lines (33,47). In the current study, MPAS scores > 0 were identified in 8 tumors. Plotted by hierarchical clusters, we found that none of the H1 PNST tumors had positive MPAS scores. In contrast, 8 tumors within the H2/H3/H4 FSA clusters had positive MPAS scores (Supplementary Figure S18). These included tumors carrying oncogenic PDGFB fusions, the SPRED1-FAM98B fusion, EGFR T724M, and HRAS G13R. The other 2 tumors with positive MPAS values had no obvious MAP kinase activators. Surprisingly, the single tumor bearing NF1 A699Ifs*3 did not have a positive MPAS score, however, the allelic frequency of this variant was only 0.1, so loss of this TSG may have been insufficient to promote measurable MAP kinase pathway activation compared to oncogenic activators. This same tumor also had a low frequency frameshift mutation in MAP3K1 G394Kfs*28. Overall, this analysis indicates that, unlike human malignant PNST, inactivation of NF1 and the resultant MAP kinase pathway activation are not characteristics of canine PNST, while some canine FSA exhibit MAP kinase activation in response to oncogenic drivers and may be amenable to treatment with targeted inhibitors.

Our RNA sequencing data validated 47% of the previously identified markers for canine FSA (4,5). This could be due to variability in analyses techniques and/or histological characterization of canine PNSTs. Similar to our study, a recent RNAseq study of canine STS subtypes, also showed elevated expression of *GLI* and *CLECB1* in PNST samples (48), but failed to confirm elevation of S100 and vimentin in PNST samples. Further studies at both transcript and protein levels are required to determine specific molecular markers for PNST.

With respect to the STS tumor micro-environment, the infiltrating CD3+ T-cell density as assessed in each tumor by IHC correlated with *in silico* quantified CD8+ T-cell abundance. Further, expression of several current actionable co-stimulatory or co-inhibitory immune checkpoint targets, including PDL1 and CTLA4, positively correlated with the number of infiltrating T-cells. These data are consistent with observations in murine pre-clinical models and human tumors wherein upregulation of inhibitory immune checkpoint molecule expression on CD8+ tumor-infiltrating lymphocytes (TILs) is a critical mechanism by which cancer cells can escape the antitumor immune response (49–51) Moreover, these data also suggest that, like human cancer patients, high T-cell infiltration of canine soft tissue sarcoma tumors as a determinant of probability of clinical response to cancer immunotherapies warrants further investigation (52,53). However, another marker of immunotherapy, tumor mutational burden, did not correlate with T cell density in this study. While several studies have shown improved responses to immune checkpoint inhibitors in tumors with high mutational burden (54,55), a recent pan-cancer study revealed no significant association between mutational burden and CD8+ T-cell infiltration; however, tumor histo-type specific associations do exist (56). Interestingly, most abundant tumor microenvironment cell type was B cells in 41% of STS canine samples. A recent gene expression profiling study of 608 human STS tumors identified B cells as the strongest prognostic factor associated with patient immunotherapy response and survival (57). A similar categorization of canine patients with STS can help classify cohorts that may positively respond to immunotherapy.

In summary, this study shows STS tumors are diverse with respect to transcriptomic, genomic and immunogenomic profiling. The H1 cluster of PNSTs, largely comprised of grade I and II tumors, had few variants, with 6 of 11 tumors having no identified putative SNV mutational drivers. However, copy number losses of *RB1* were observed in 4 of the 11 H1 tumors. The H2 cluster was notable for the presence of *PDGFB* gene fusions in 3 of the 7 tumors. Of the 6 tumors bearing *TP53* driver mutations, 50% were from cluster H3 (n=5). The H4 cluster exhibited elevated immune activation, T-cell infiltration, and a lack of clock-like COSMIC signatures. MAP kinase pathway activation was only observed in clusters H2-H4 with 90% of the grade III tumors. Although prognosis for this cancer is improved by surgical intervention, overall survival of dogs with grade III tumors is poor compared to dogs with grade I tumors (58). Thus, the molecular and genetic driver characterization of canine STS subtypes may aid in diagnosis and the identification of novel therapies that improve the outcomes for canine and human patients with recurrent or metastatic disease.

## Acknowledgements

We would like to thank Irene Mok and Lindsay Carroll for Tissue Archive curation. We acknowledge the help of Marina Schlaepfer and Rujuta Idate in screening variants for putative driver mutations.

