## Supplementary Figures for "Integrated analysis of canine soft tissue sarcomas identifies recurrent mutations in *TP53, KMT* genes and *PDGFB* fusions"

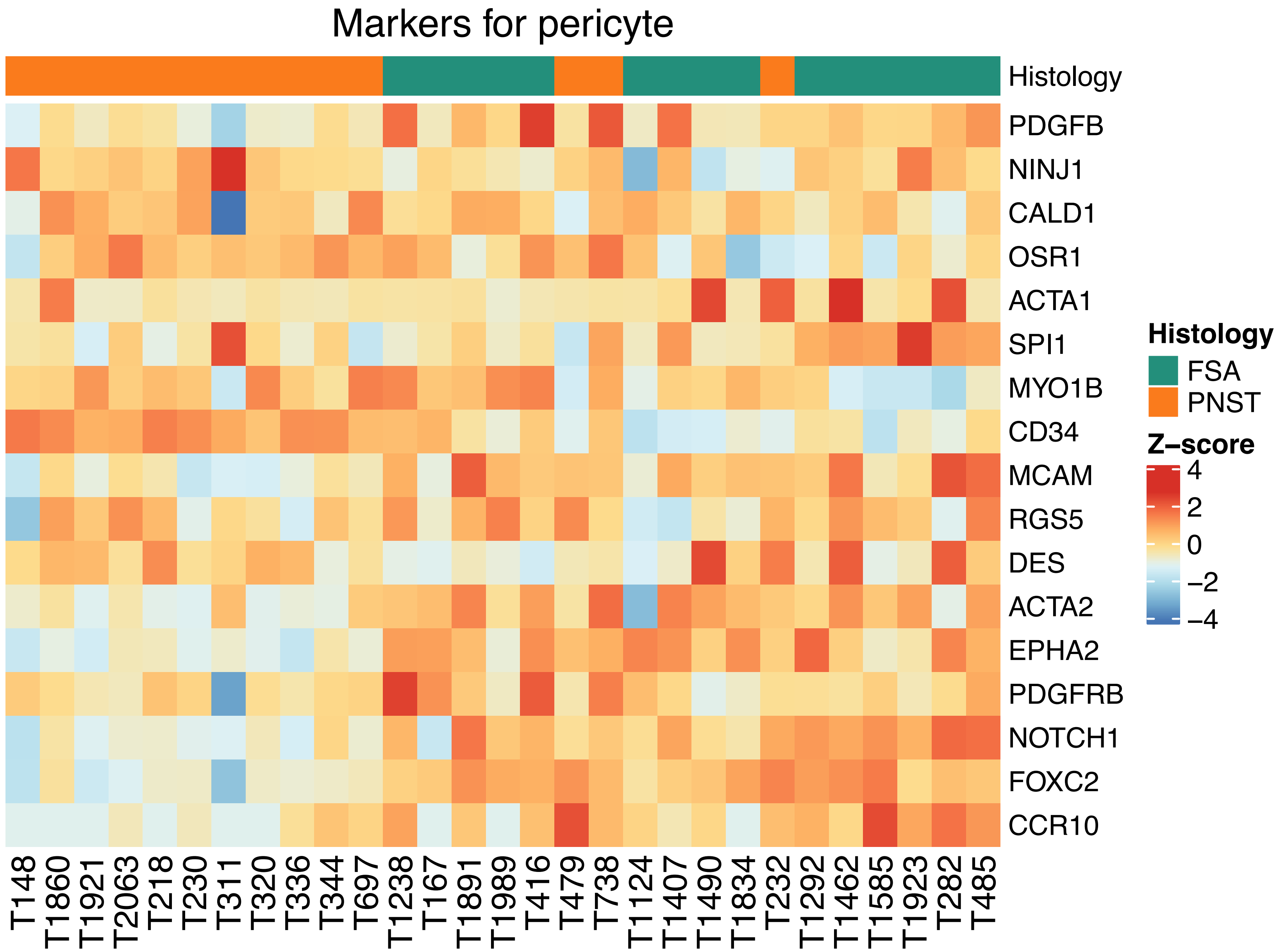


**Supplementary Figure S1**. Heatmap of pericyte marker genes. Expression levels of 17 genes identified as pericyte markers in the literature and gene ontology databases GO:1990874 and GO:1904238 are plotted.

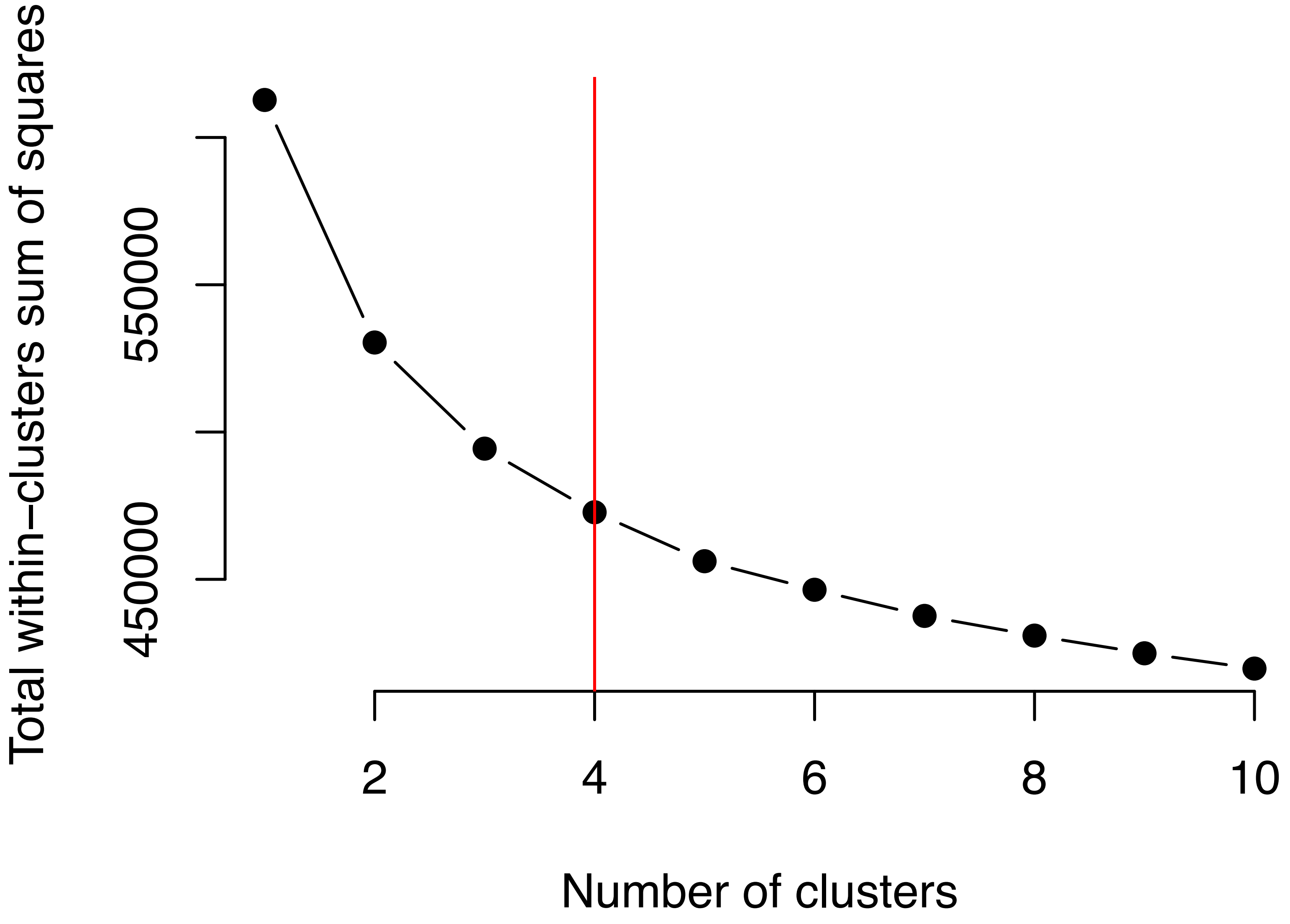


**Supplementary Figure S2.** The Elbow method was used to determine the optimal number of clusters for 29 soft tissue sarcoma tumor samples.

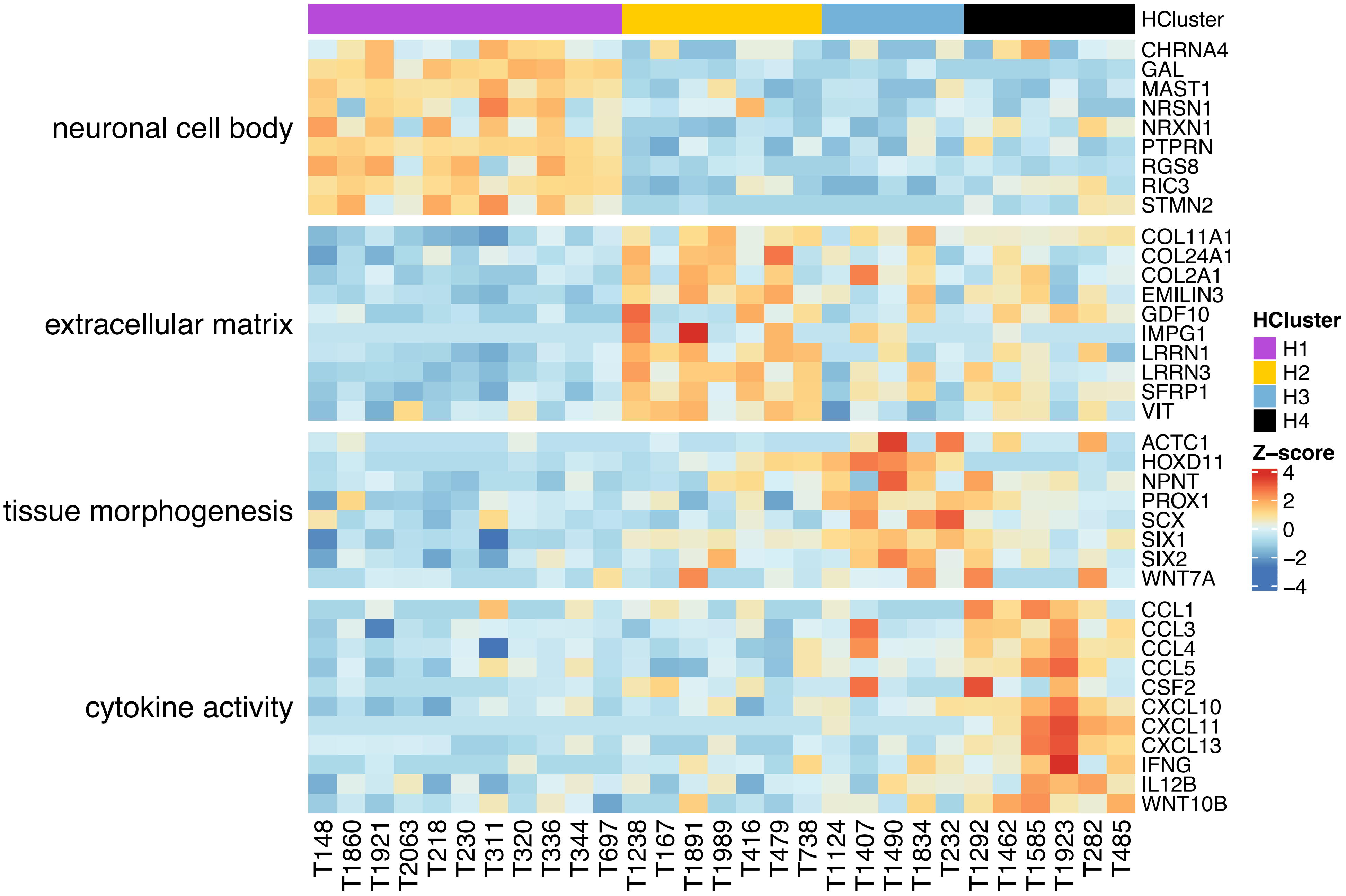


**Supplementary Figure S3.** Heatmap of differentially expressed genes in selected enriched pathways within each of the four clusters.

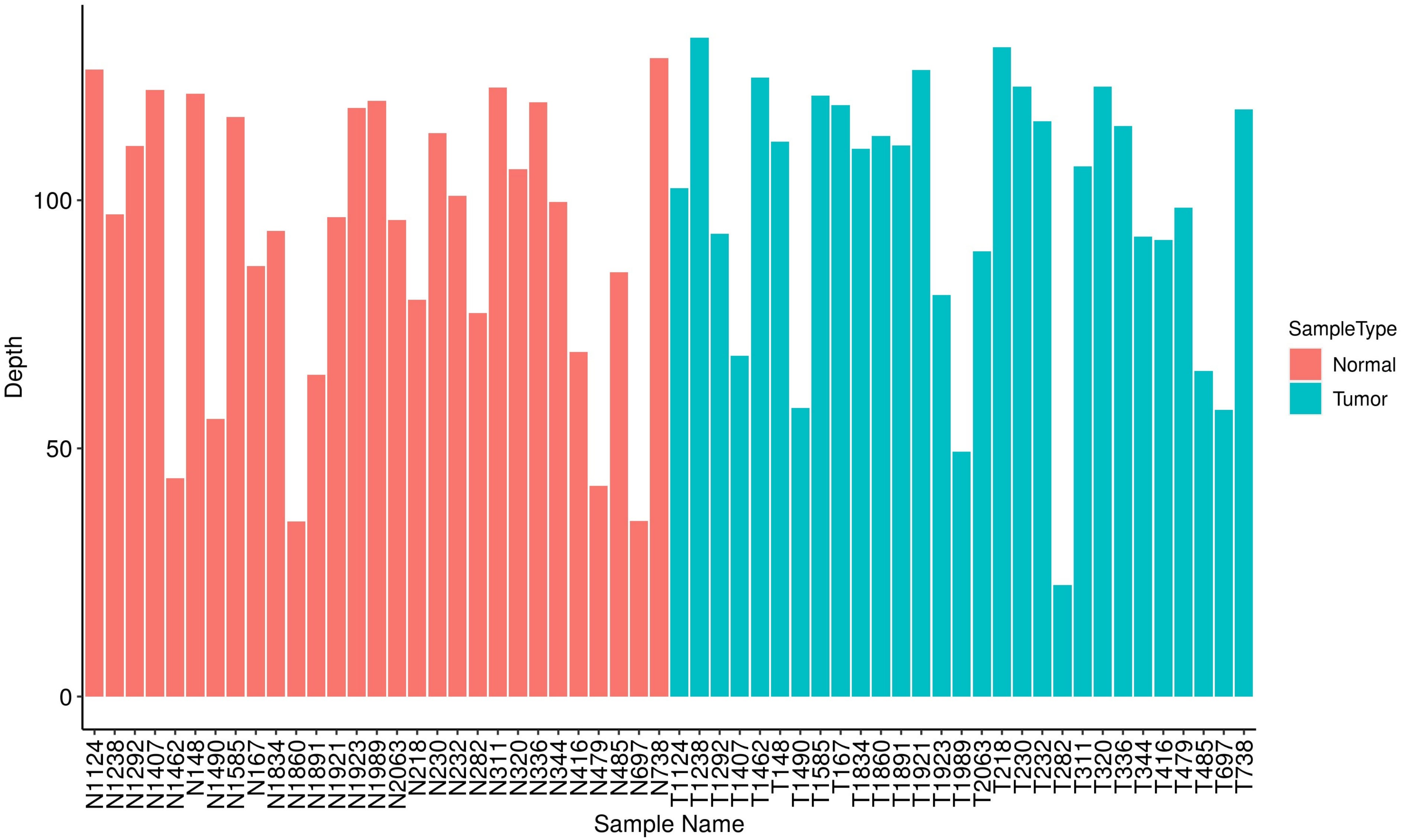


**Supplementary Figure S4.** Whole exome sequence mapping coverage/depth across 29 soft tissue sarcoma tumors and their matched normal samples.

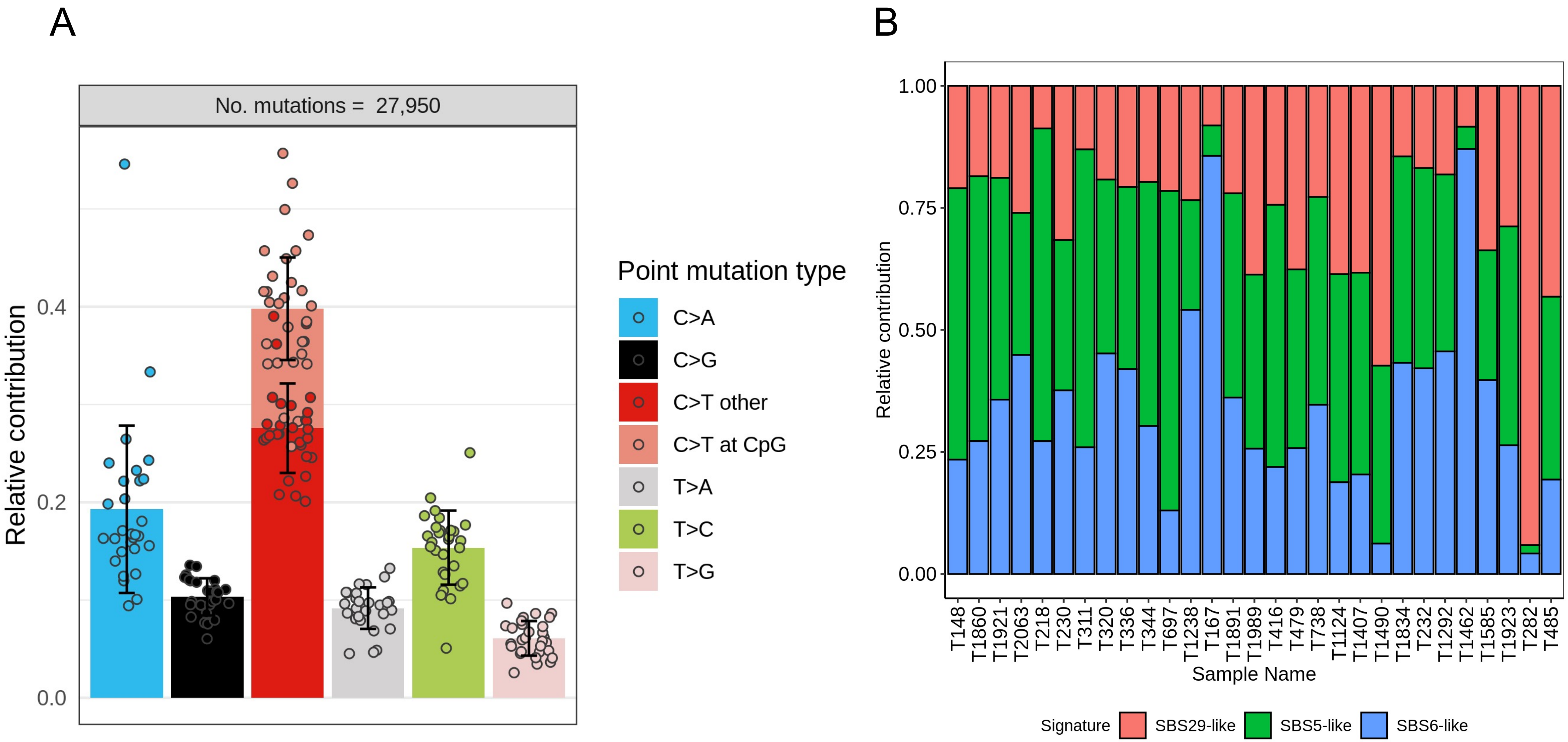


**Supplementary Figure S5.** Mutational signature identification. A. Relative contribution of 6 single nucleotide substitutions in 29 STS samples. B. Relative contribution of three *de-novo* mutational signatures in each of the 29 STS tumors.

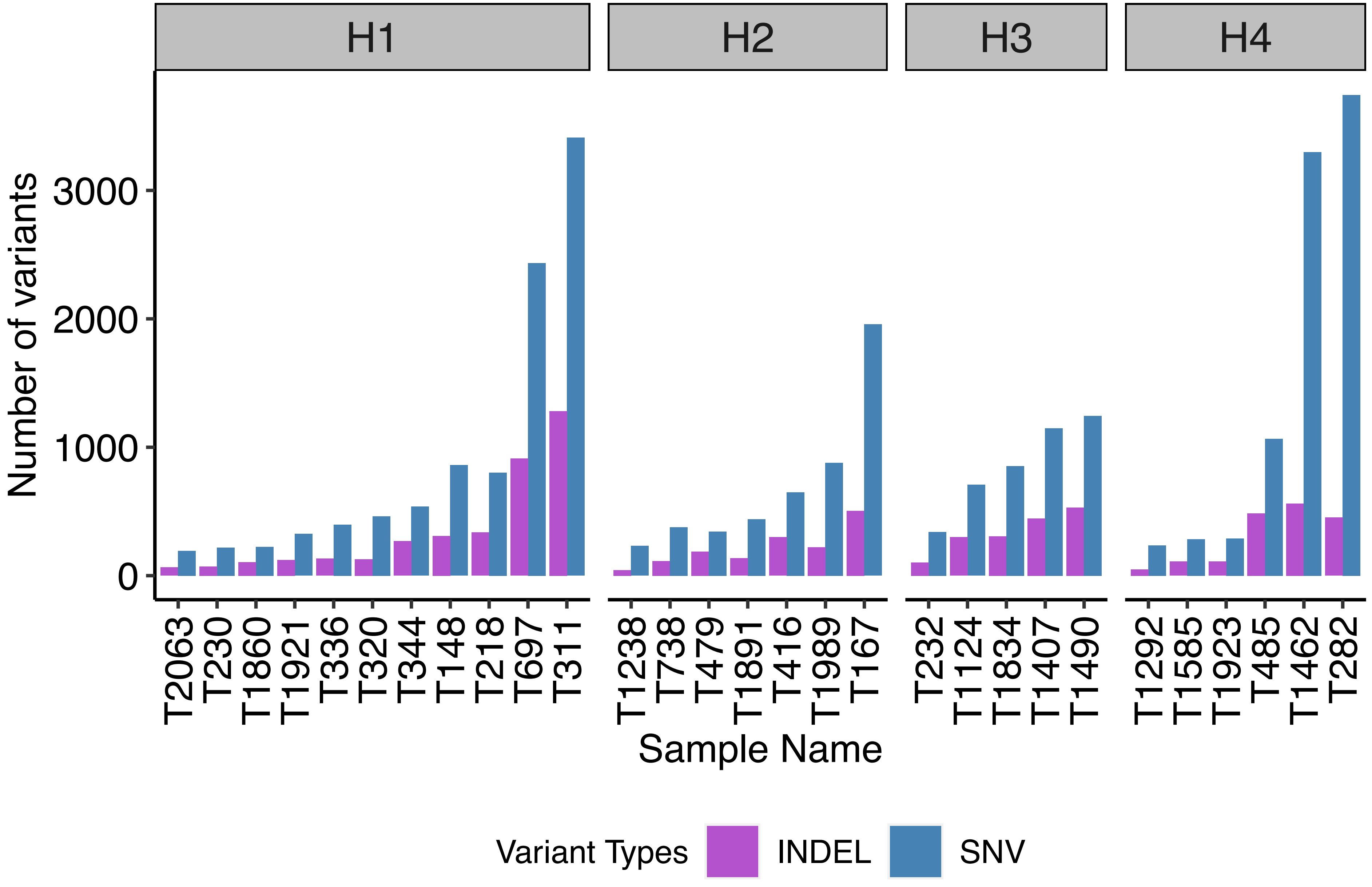


**Supplementary Figure S6.** Number of non-synonymous somatic SNV and INDELs identified by Mutect2. The 29 soft tissue sarcoma tumor samples are arranged by their *in silico* generated cluster ID.

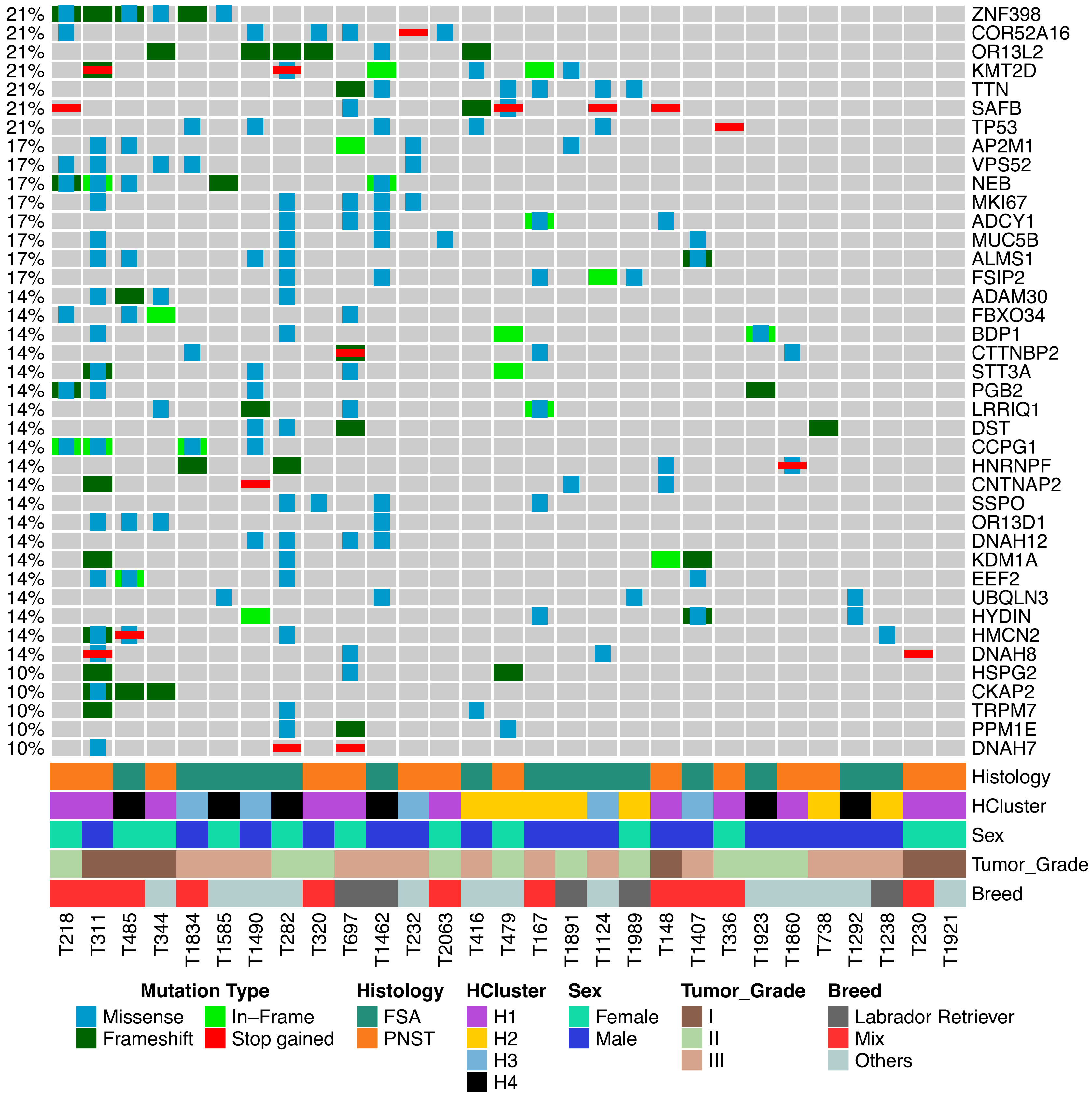


**Supplementary Figure S7.** Oncoplot of top 40 recurrently mutated protein coding genes. The samples are arranged based on mutual exclusivity of the gene mutations across 29 soft tissue sarcoma samples.

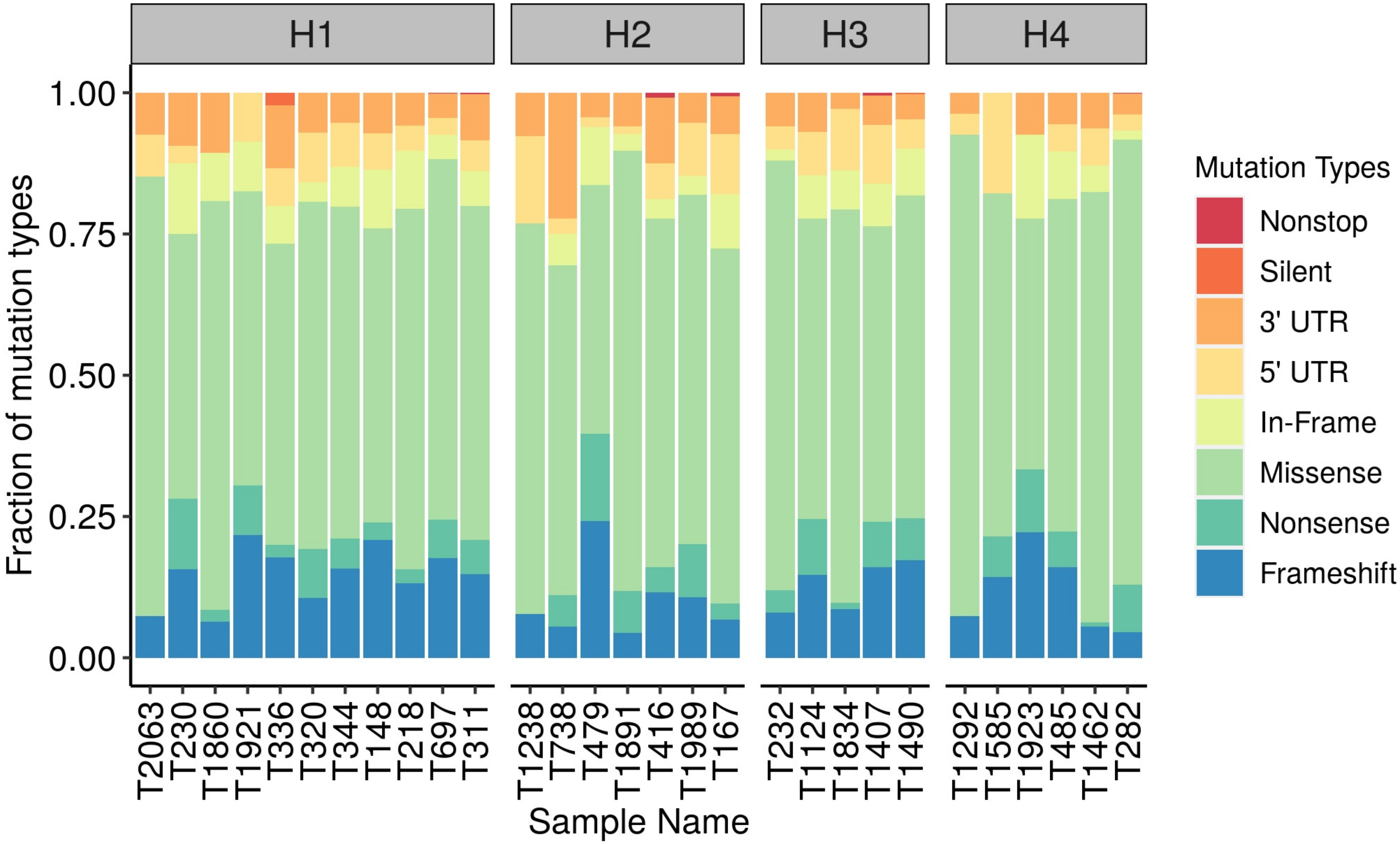


**Supplementary Figure S8.** Distribution of mutation types across 29 canine soft tissue samples.

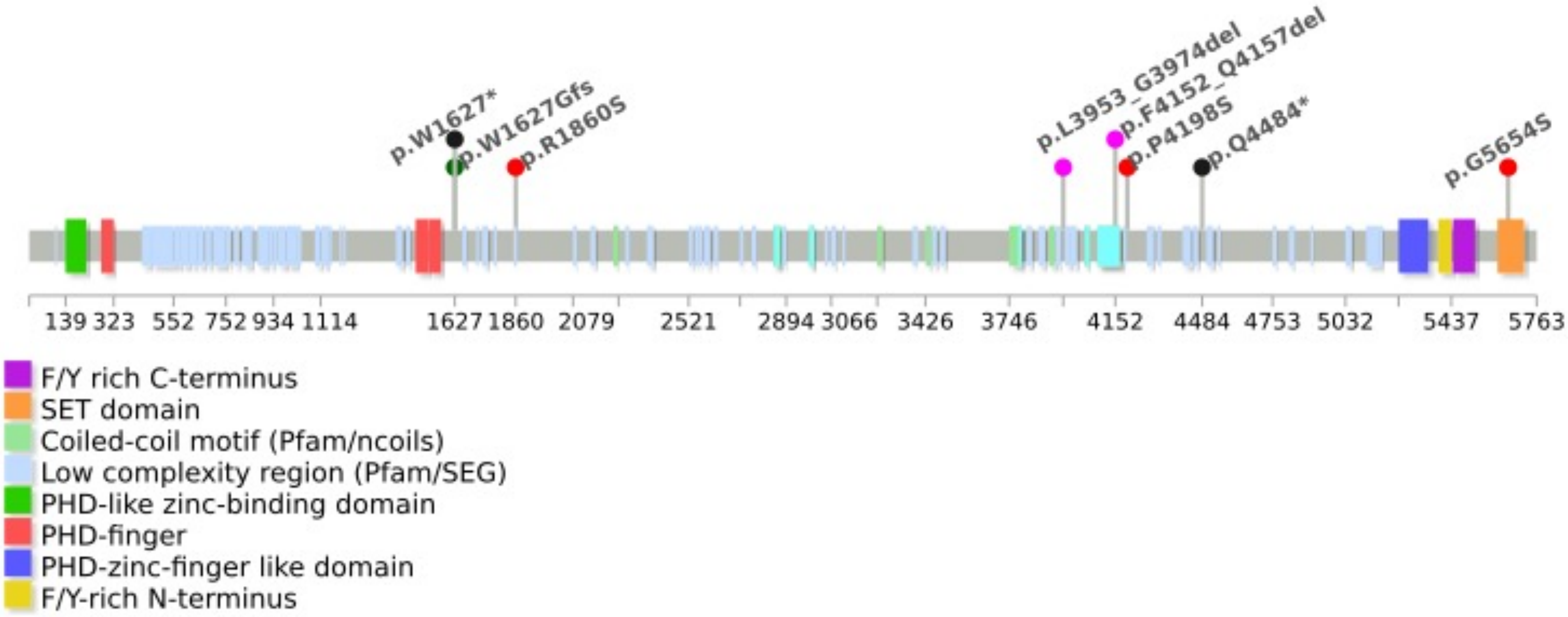


**Supplementary Figure S9.** Lollipop plot of KMT2D protein showing distribution of non-synonymous mutations identified in 21% of canine soft tissue sarcoma samples.

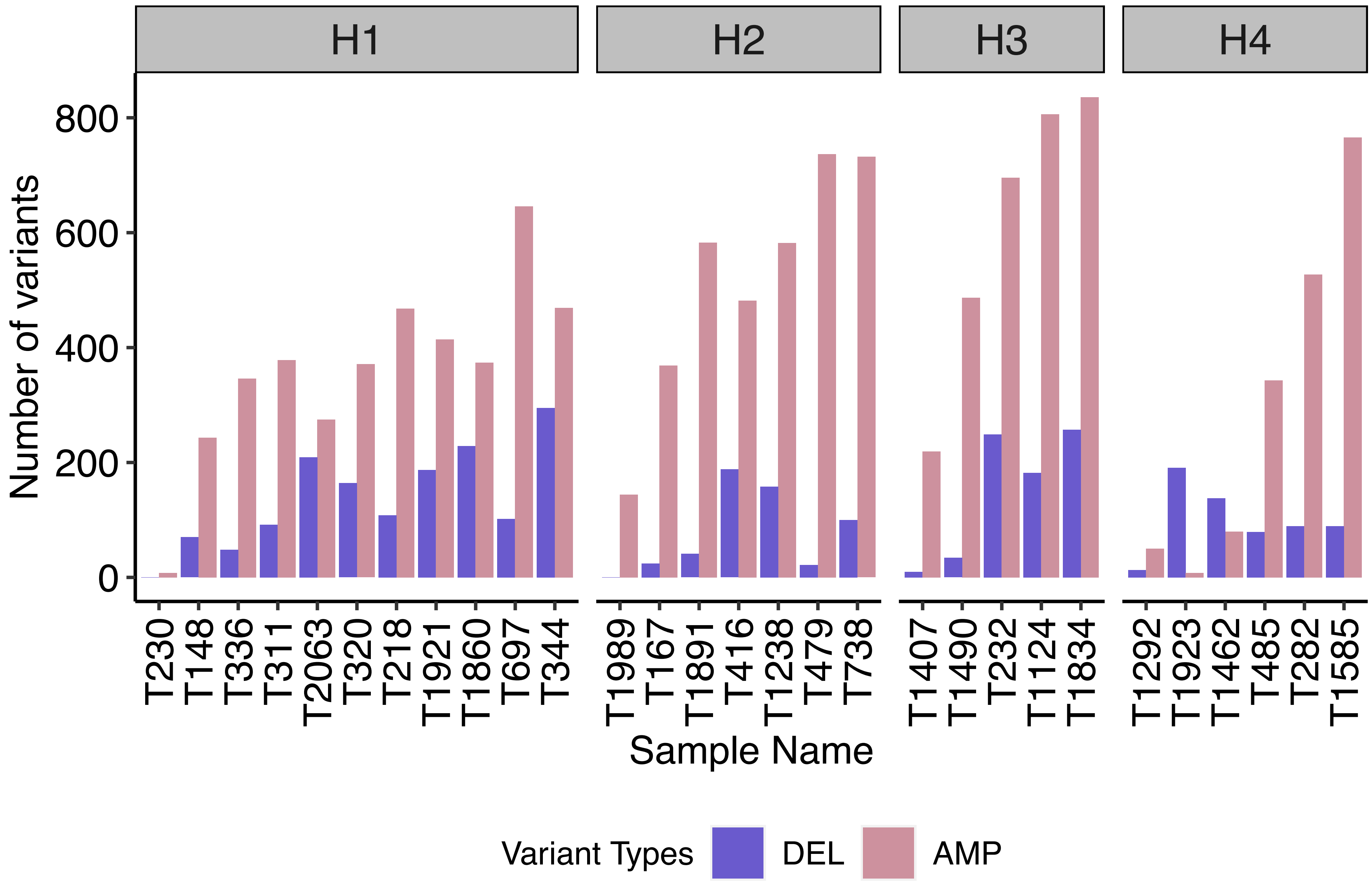


**Supplementary Figure S10.** Number of genes within significantly amplified or deleted regions of the genome as identified in 29 soft tissue sarcoma samples.

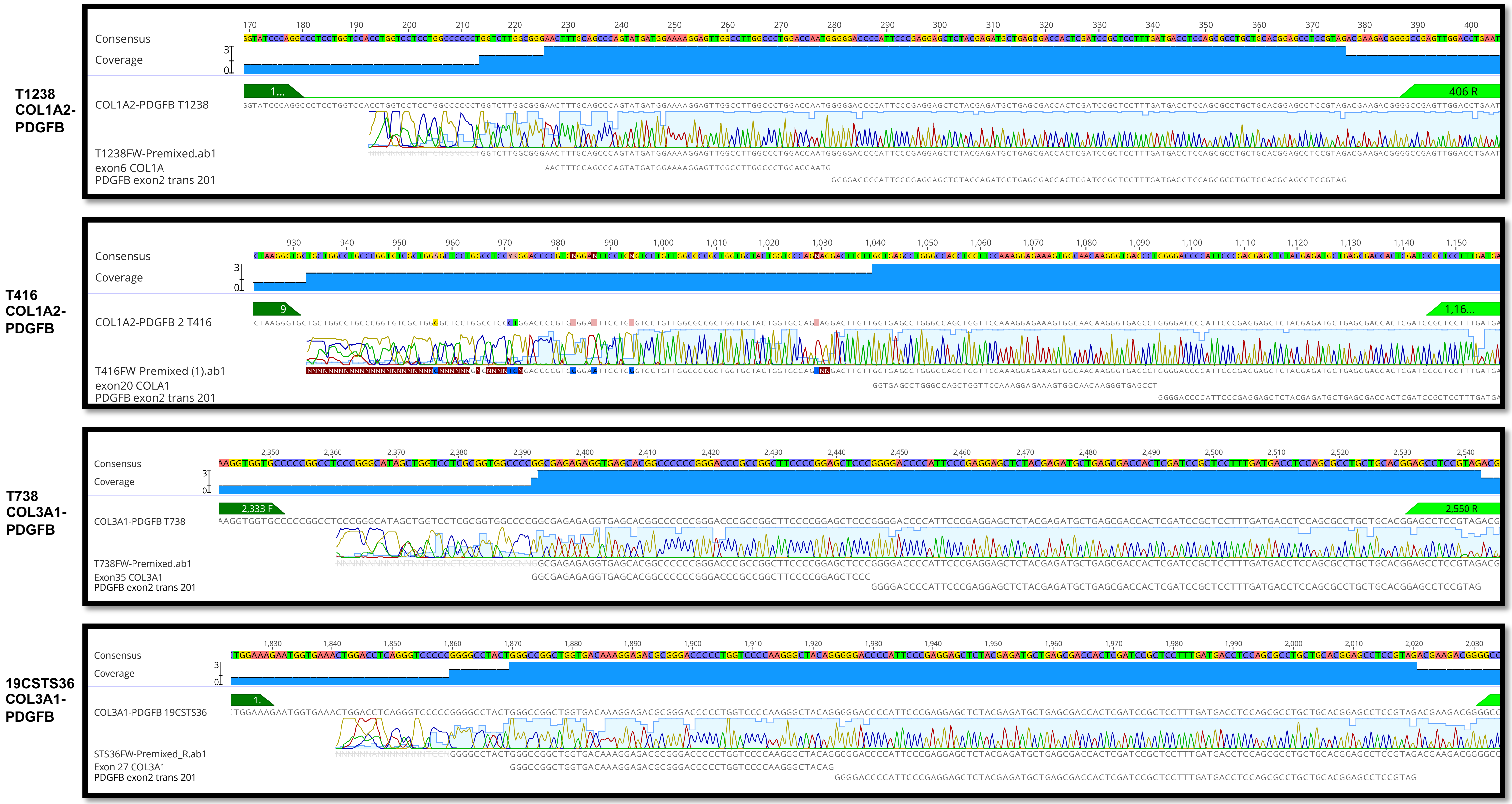


**Supplementary Figure S11**. Validation of fusion genes via Sanger sequencing. Total RNA extracted from tumors was reverse transcribed and cDNA was amplified using primers designed to span the gene fusion breakpoint. Amplified regions were sequenced and aligned to the fusion reads, as well as the respective sequences of the fused exons.

| Fusion | Forward primer | Reverse Primer |
|---|---|---|
| T1238 COL1A2-PDGFB | 5’-GTGACGATGGTATCCCAGGC | 5’-AATTCAGGTCCAACTCGGCC |
| T416 COL1A2-PDGFB | 5’-CCTGACTGGTGCTAAGGGTG | 5’-TGGAGGTCATCAAAGGAGCG |
| T738 COL3A1-PDGFB | 5’-ATAAGGGTGAAGGTGGTGCC | 5’-GTCTTCGTCTACGGAGGCTC |
| FACC-19-CSTS36 COL3A1-PDGFB | 5’-CTCAGGGTCCTGCTGGAAAG | 5’-AATTCAGGTCCAACTCGGCC |

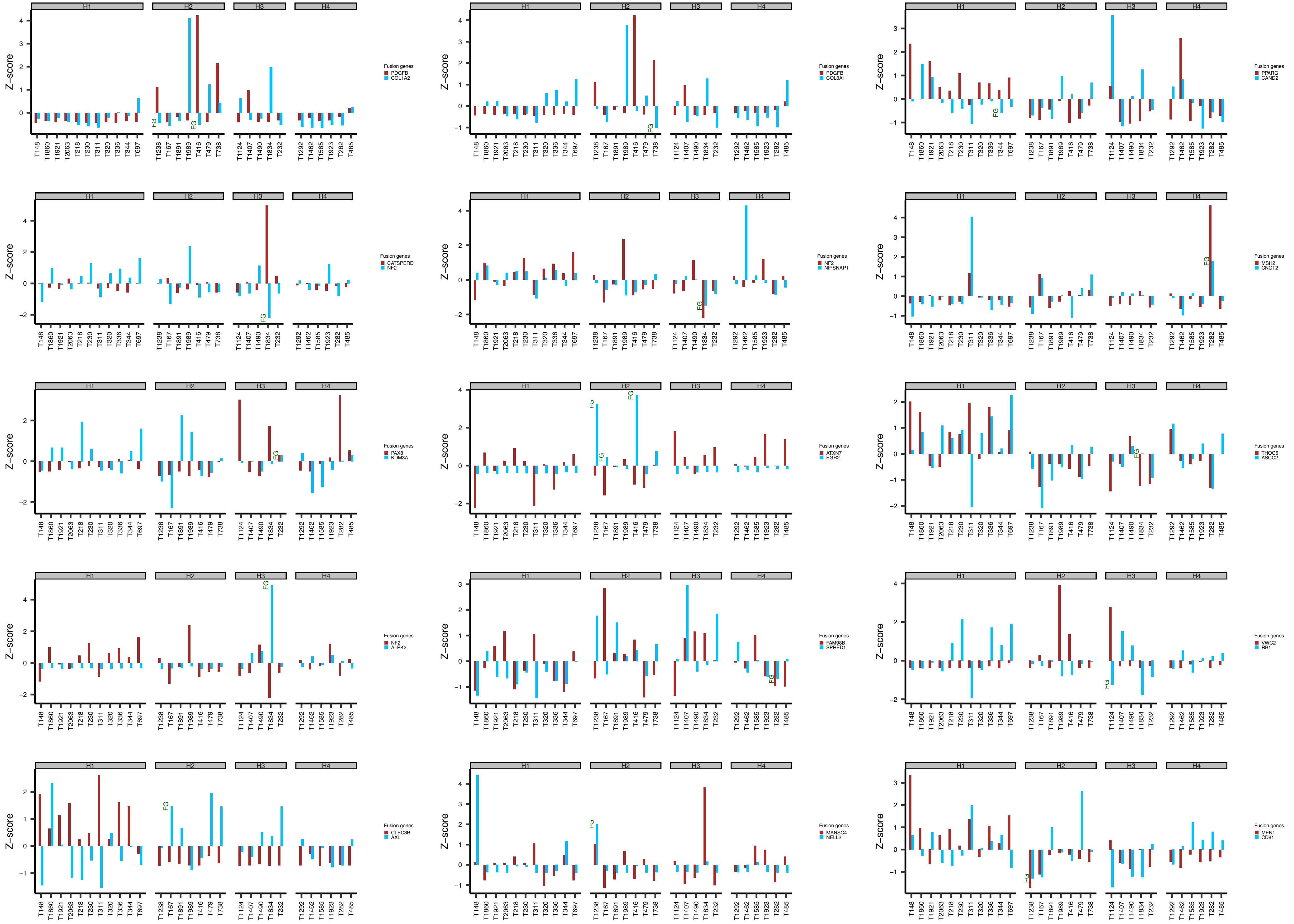


**Supplementary Figure S12**. Gene expression plots of fusion gene partners across 29 soft tissue sarcoma samples. The samples with identified fusions are labeled FG to the right of the gene partners in each sub-plot.

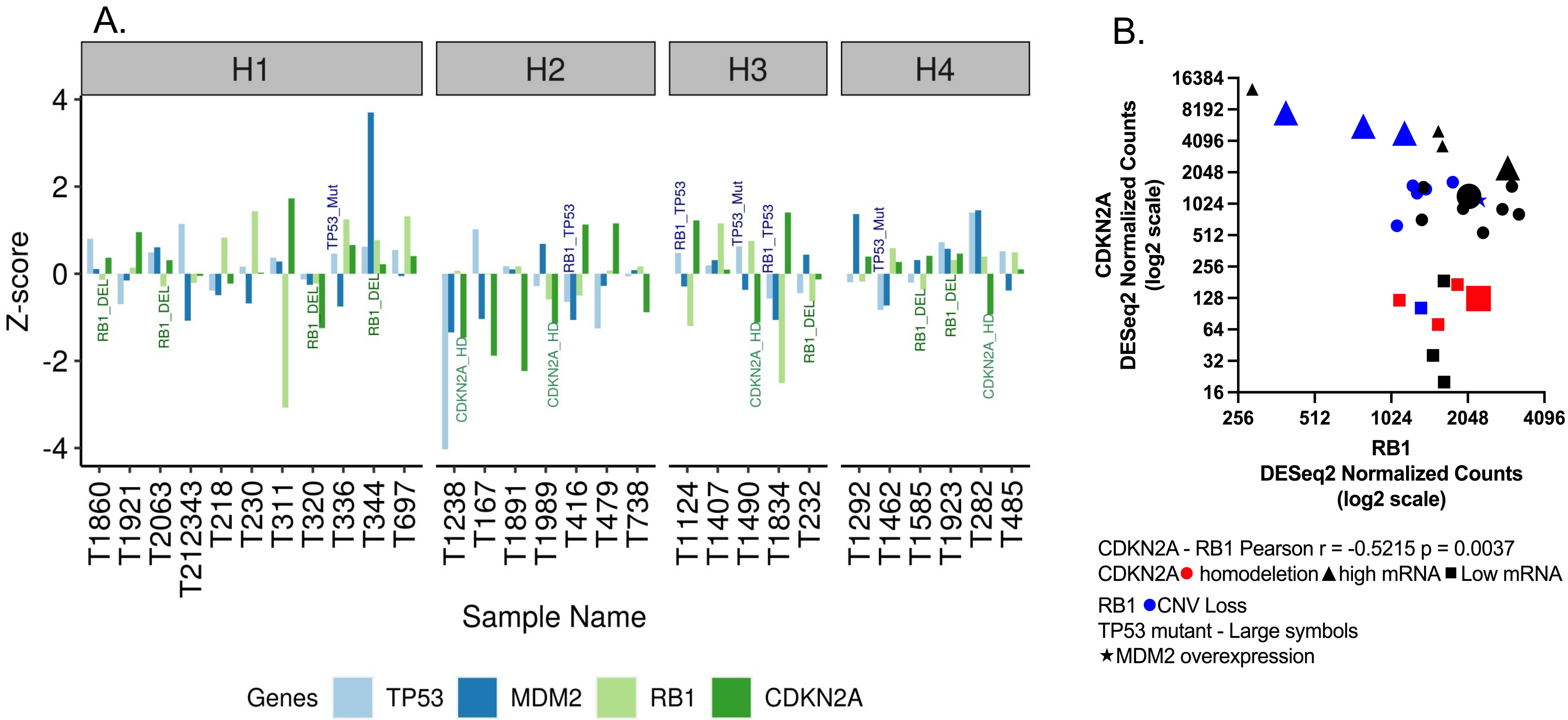


**Supplementary Figure S13.** Expression levels of cell cycle related genes. A. Z-scores of 4 genes are plotted across 29 samples in 4 clusters. The copy number deletions are denoted by DEL, homozygous deletions by HD, and SNV/INDEL mutations by MUT. B. Correlation between *CDKN2A* and *RB1* gene expression. There was a significant negative correlation between the expression levels of *CDKN2A* and *RB1*. Other identified gene deletions or expression changes are marked as indicated in the symbol legend.

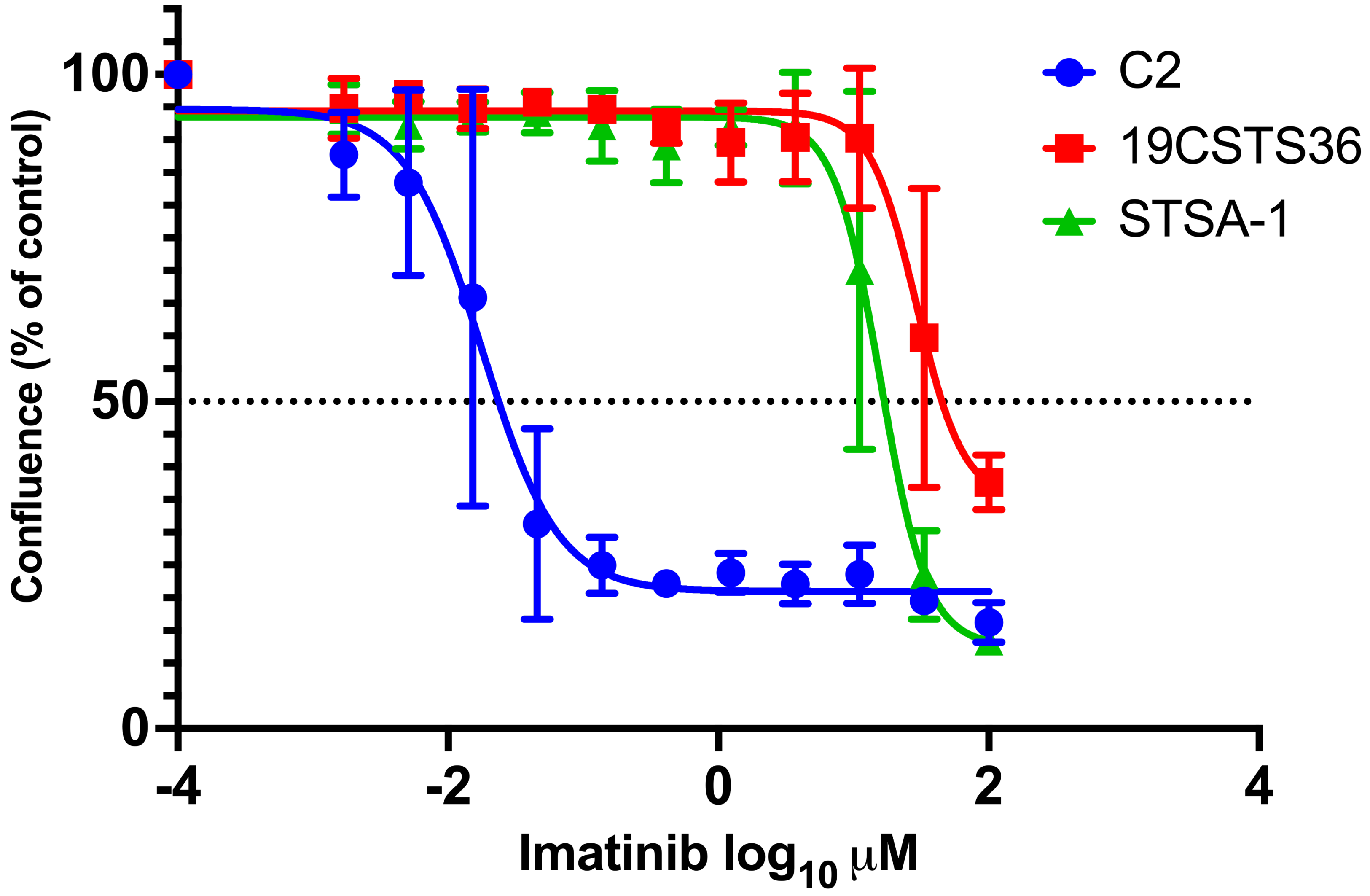


**Supplementary Figure S14. Sensitivity of canine cell lines to Imatinib, a multi-target inhibitor of PDGFR, v-Abl, and c-Kit.** Cells were plated on 96 well plates and treated with serial dilutions of imatinib mesylate 24 hours later. After treatment for 72 hours, cell confluence was measured, normalized to time zero, and expressed as percent of control. Data symbols represent mean ± SD of 3 experiments with averaged triplicate wells. Non-linear curves (4 parameter) of log dose versus percent of control were fit and LD50 values were interpolated as the dose at which cell number was 50% of control.

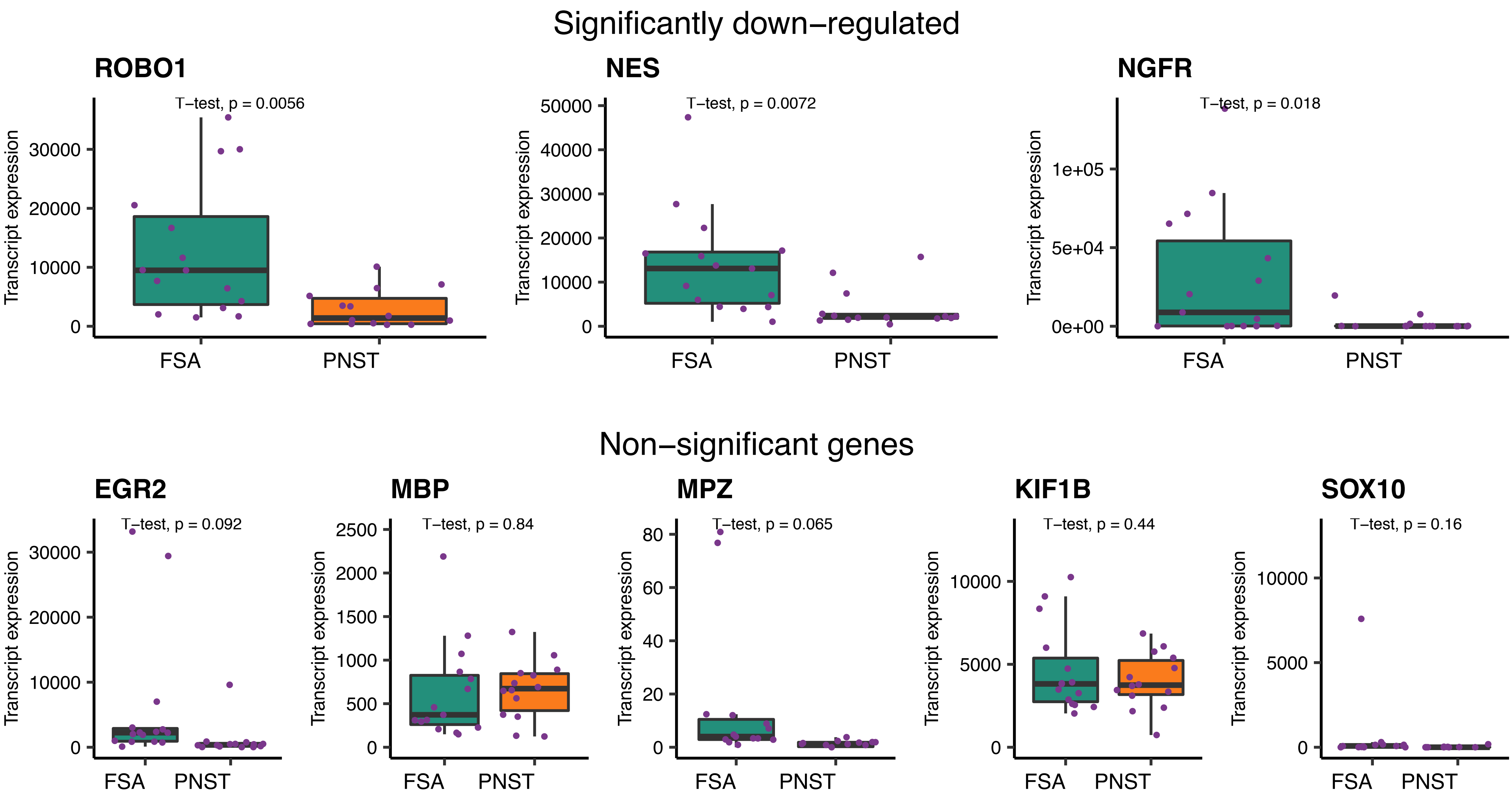


**Supplementary Figure S15.** Expression of previously identified PNST markers. Unlike the previously reported data, these genes were either significantly down-regulated or were not variably expressed when compared to their expression in FSA samples.

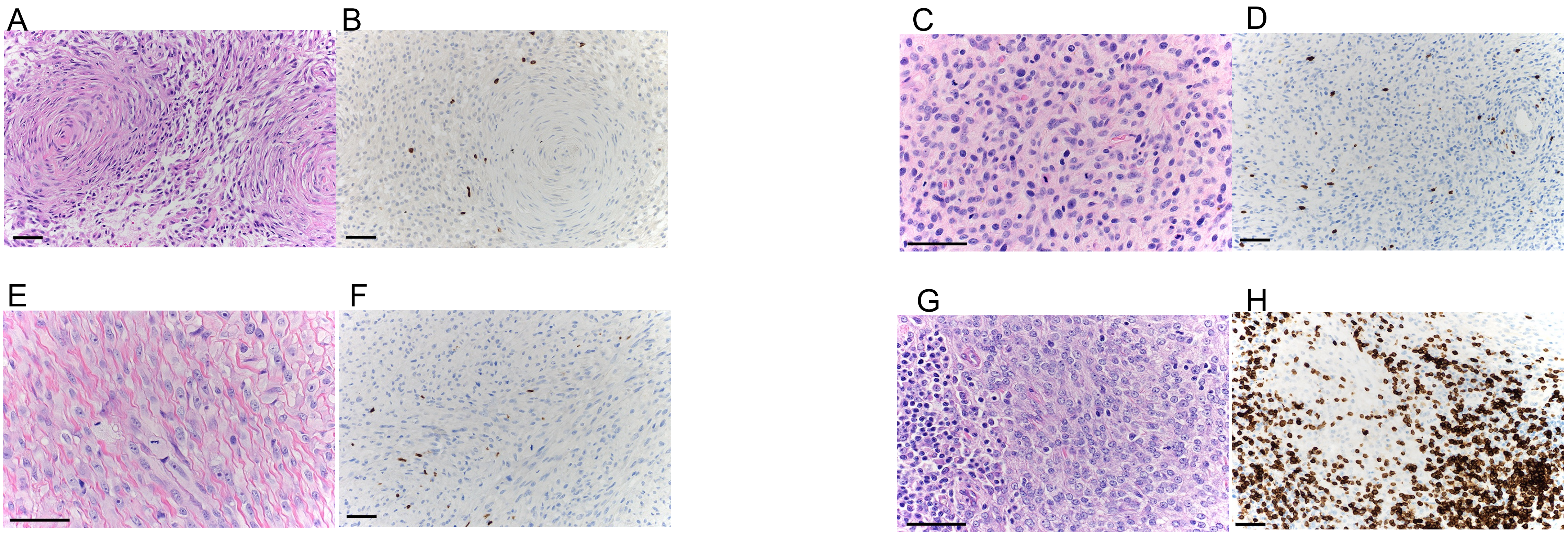


**Supplementary Figure S16.** Representative histological images from each cluster. **A.** H&E image from T336 (H1 cluster) demonstrating typical histomorphology of a malignant peripheral nerve sheath tumor with characteristic concentric whirling of neoplastic spindle cells around central collagen. 20x magnification, scale bar = 50μM. **B.** Corresponding image of immunohistochemically labeled CD3+ tumor-infiltrating leukocytes. 20x magnification, scale bar = 50μM. DAB chromogen, hematoxylin counterstain. **C.** H&E image from T1238 (H2 cluster) demonstrating tumor cells with relatively uniform nuclei with indistinct nucleoli, scant cytoplasm, and are embedded within abundant collagenous stroma. H& E with 40x magnification, scale bar = 50μM. **D.** Corresponding image of immunohistochemically labeled CD3+ tumor-infiltrating leukocytes. 20x magnification, scale bar = 50μM. DAB chromogen, hematoxylin counterstain. **E.** Neoplastic cells in T1124 (H3 cluster) display moderate nuclear and cellular pleomorphism, with large round to oval nuclei, prominent nucleoli, occasional giant cells, and increased mitotic figures. Cells are embedded in a prominent collagenous stroma. H&E. 40x magnification, scale bar = 50μM. **F.** Corresponding image of immunohistochemically labeled CD3+ tumor-infiltrating leukocytes. 20x magnification, scale bar = 50μM. DAB chromogen, hematoxylin counterstain. **G.** Neoplastic cells in T1585 (H4 cluster) are more anaplastic and display moderate nuclear pleomorphism with round to oval nuclei containing 1-2 prominent nucleoli, scant cytoplasm, and indistinct cell borders. A peripheral aggregate of infiltrating lymphocytes is observed. H&E. 40x magnification, scale bar = 50μM. **H.** Corresponding image of immunohistochemically labeled CD3+ tumor-infiltrating leukocytes. 20x magnification, scale bar = 50μM. DAB chromogen, hematoxylin counterstain.

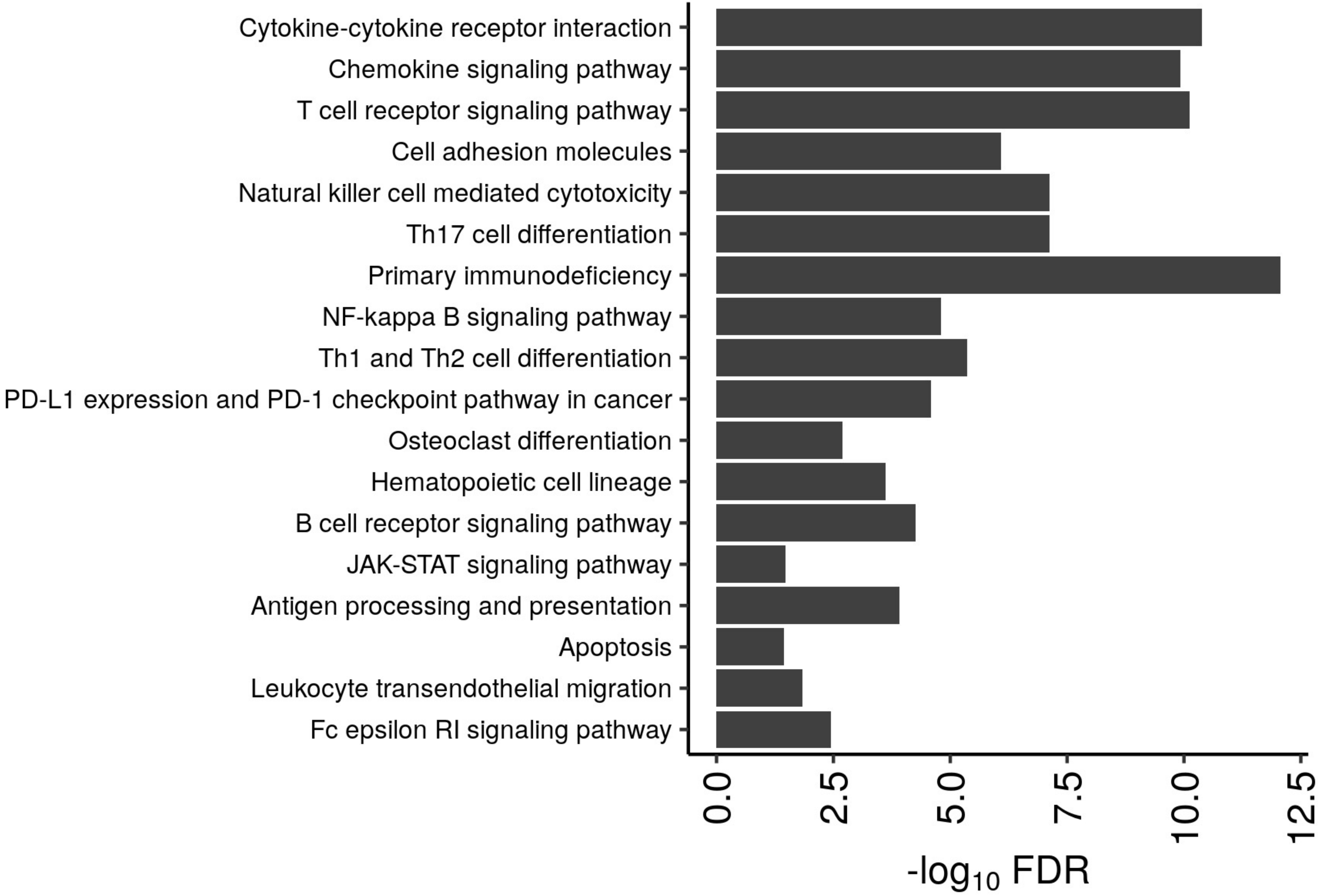


**Supplementary Figure S17.** Enriched KEGG pathways that are significantly correlated with CD3+ T cell infiltration in tumors as quantified by IHC.

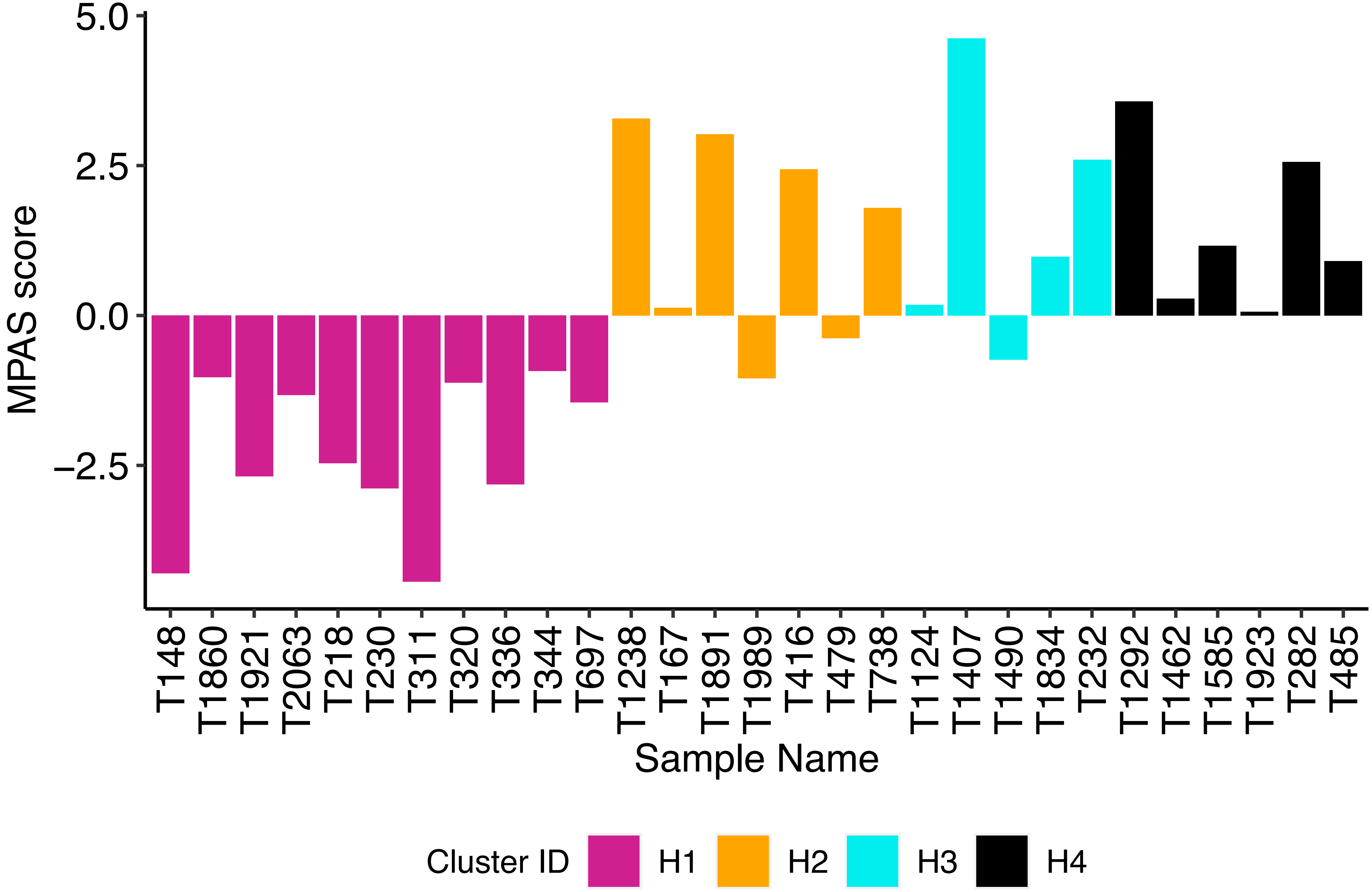


**Supplementary Fig. S18.** MAP Kinase Pathway Activation score (MPAS) of 29 soft tissue sarcoma samples arranged by their respective clusters.
