## Supplementary Methods for "Integrated analysis of canine soft tissue sarcomas identifies recurrent mutations in *TP53, KMT* genes and *PDGFB* fusions"

### Tumor sample selection and categorization

Archived primary tumor samples with a diagnosis of soft tissue sarcoma were obtained from the Colorado State University (CSU) Flint Animal Cancer Center's tissue archive. Tumor tissues are archived from canine patients presenting to the Flint Animal Cancer Center in compliance with the CSU Institutional Animal Care and Use Committee following obtainment of owner consent. Tumors samples were reviewed by a board-certified veterinary anatomic pathologist (DR) and selected based on 1) quality of the formalin-fixed paraffin embedded tissue, assessed by hematoxylin and eosin (H&E)-stained slide evaluation confirming adequate overall biopsy size and a high degree of viable tumor tissue, 2) availability of a corresponding fresh frozen tumor biopsy sample for RNA extraction, and 3) availability of normal tissue or whole blood for isolation of normal genomic DNA. Tumors were histologically graded as previously described (1,2) and given a diagnosis corresponding to one of two broad histologic subtypes based on patterns of cell growth: 1) fibrosarcoma, based on the presence of a solid or fasciculated growth pattern, or 2) malignant nerve sheath tumor/perivascular wall tumor, based on the predominance of any mixture of palisading, Antoni A or Antoni B, or concentric whorling growth patterns. Grading and diagnosis was performed in a blinded fashion without any a priori knowledge of transcriptomic clustering.

**Pericyte marker genes**

In order to distinguish between PWT and PNST at a molecular level, we used literature searches to catalog genes that are used as markers for pericytes (3-5). We also used the gene ontology terms, vascular associated smooth muscle cell proliferation (GO:1990874) and pericyte cell differentiation (GO:1904238) to curate additional markers for pericytes.

**Clustering tumor samples**

Unsupervised hierarchical clustering was used to group 29 soft tissue sarcomas into distinct clusters. A subset of genes (n=1,926) with mean expression $\log_2 > 2$ and mean variance $\log_2 > 5$ was used to cluster the tumor samples. The optimal number of hierarchical clusters was determined using the elbow method (6). Spearman correlation and Ward's minimum variance method (Ward.D2) was used to calculate distance and cluster STS samples, respectively.

**Fusion gene identification**

A gene fusion database was created from CanFam3.1 genome assembly and ensembl gene annotation (version 99) by using prep_genome_lib.pl script from Star-Fusion (7). The gene fusions were identified by STAR-fusion tool where trimmed fastq reads were used as input. The fusions were manually curated to identify putative drivers. Fusion genes in which both partners were part of same gene family (e.g. CUX1--CUX2) and genes with no known functions and/or without gene symbols were eliminated. The fusion genes were further curated to identify fusions with known cancer genes (OncoKB database). Selected fusion genes were also validated via Sanger sequencing. The primers were designed from the STAR-fusion predicted cDNA sequence of fusion genes using Geneious Prime 2021.2 software (https://www.geneious.com). The forward and reverser primer sequences were reported in Supplementary Figure S11.

**Differential expression**

We ran four sets of pair-wise comparisons to identify differentially expressed genes (DEGs) between samples from each cluster and combined samples from the other clusters (e.g. H2 vs H1, H3, H4) using DESeq2 in R environment (8). Briefly, the raw counts for all genes were filtered to retain genes with at least 4 reads in at least 4 samples which resulted in 18,201 genes that were processed for downstream analysis. Following normalization of raw counts, the DEGs were identified using DESeq function. The DEGs with fold change of $\log_2 > 3$ and adjusted p-value $< 0.05$ were used as input for identifying pathways that were up-regulated in each cluster. The functional annotation of genes was done using g:Profiler R package and sources included KEGG, Reactome, WikiPathways, and Gene Ontology databases (9). The pathways with false discovery rate of <0.05 were considered enriched.

**Somatic simple variant annotation**

The VCF files with somatic variants were converted to MAF (Mutation Annotation File) format (https://docs.gdc.cancer.gov/Data/File_Formats/MAF_Format/) using the perl code: vcf2maf.pl (https://github.com/mskcc/vcf2maf). Each variant was mapped to only one of all possible gene transcripts using the "canonical" isoform from Ensembl database (v99).

**Homozygous deletion identification**

Genes with homodeletion in 29 STS tumor samples (Supplementary Figure S13), were identified using VarScan2 from the whole exome sequence data (10). The candidate homozygous deletions were called using *copyCaller* function and by specifying the option *--output-homdel-file*.

**Data plotting**

The oncoplots and heatmaps were plotted using ComplexHeatmap R package (11). Barcharts and boxplots were plotted using R package ggplot2 (12). The lollipop plots were created by Lollipops software using canine uniport ID (13).

**Immunohistochemical analysis**

H&E-stained slides were evaluated by a board-certified pathologist (DPR) to confirm diagnosis and the presence of adequate viable tumor tissue for analysis. Immunolabeling was performed via routine, automated methods on the Leica Bond Max autostainer (Leica Biosystems Inc.) as previously described (14), using a mouse monoclonal anti-human CD3 (pan T lymphocyte marker; Leica, clone LN10, ready-to-use format). Antigen retrieval was performed using Leica Epitope Retrieval 2 (Tris-EDTA buffer, pH 9) for 20 min. Primary antibody incubation was carried out at room temperature for 30 min, followed by detection with PowerVision IHC detection systems (Leica Biosystems, Inc.), using a polymeric horseradish peroxidase anti-mouse IgG and DAB chromogen. Slides were counterstained with hematoxylin.

Whole slide images of IHC stained slides were digitally captured using an Olympus IX83 microscope at 10x magnification and fixed exposure times for all samples. Quantitative image analysis was performed using ImageJ software (National Institutes of Health, NIH) as previously described (15). Briefly, whole slide images were converted to gray scale, and tumor tissue regions-of-interest were segmented from adjacent normal tissue or section artifacts by manual outlining in ImageJ by a veterinary pathologist. Positively labeled CD3+ T cells were counted using the ImageJ color deconvolution plugin (Fast Red, Fast Blue, and DAB vectors), with a minimum positive particle size threshold of 10 pixels and a positive pixel threshold for all immune cell markers set based on two standard deviations above the mean of appropriate isotype-stained control slides and confirmed visually by a veterinary pathologist. Images were subjected to global, automated application of this intensity and size threshold to all images. Following image analysis, positive pixel masks of each image were evaluated by a pathologist to ensure thresholding and counting accuracy. Data was analyzed and the number of infiltrating immune cells was expressed as total positive cells per $mm^2$ of tumor tissue area.

**MPAS score assessment**

The MAPK Pathway Activation Score (MPAS) quantifies relative MAPK activity using gene expression data. The expression of 10 downstream MAPK pathway targets (CCND1, DUSP4, DUSP6 EPHA2, EPHA4, ETV4, ETV5, PHLDA1, SPRY2, SPRY4) were used to calculate the score MPAS = (sum of z-score normalized expression for MPAS genes) / ($\sqrt{10}$). MPAS score has been shown to correlate with sensitivity to MAPK inhibitors in both cancer cell lines and cancer patients (16).
